# A Novel Mouse Model that Recapitulates the Heterogeneity of Human Triple Negative Breast Cancer

**DOI:** 10.1101/2022.10.07.511231

**Authors:** Zinab O. Doha, Xiaoyan Wang, Nicholas Calistri, Jennifer Eng, Colin J. Daniel, Luke Ternes, Eun Na Kim, Carl Pelz, Michael Munks, Courtney Betts, Nell Kirchberger, Sunjong Kwon, Elmar Bucher, Xi Li, Trent Waugh, Jennifer A. Pietenpol, Melinda E. Sanders, Gordon Mills, Koei Chin, Young Hwan Chang, Lisa M. Coussens, Joe W. Gray, Laura M. Heiser, Rosalie C. Sears

**Affiliations:** Department of Molecular and Medical Genetics, Oregon Health & Science University, USA; Department of medical laboratory technology, Taibah University, Saudi Arabia; Department of Biomedical Engineering, Oregon Health & Science University, USA; OHSU Center for Spatial Systems Biomedicine, Oregon Health & Science University, USA; Brenden-Colson Center for Pancreatic Care, Oregon Health & Science University, USA; Department of Molecular Microbiology and Immunology, Oregon Health and Science University, USA; Department of Cell, Developmental & Cancer Biology, Oregon Health and Science University, USA; Division of Oncologic Sciences, Oregon Health and Science University, USA; Department of Biochemistry, Vanderbilt University Medical Center, USA; Vanderbilt-Ingram Cancer Center, Vanderbilt University Medical Center, USA; Department of Pathology, Microbiology, and Immunology, Vanderbilt University Medical Center, USA; Knight Cancer Institute, Oregon Health & Science University, USA

**Author notes:** Contributed equally (Zinab O. Doha, Xiaoyan Wang). **Additional information** - Funding for these studies include U54 CA209988, R01 CA186241, R01 CA129040, U01 CA224012, Prospect Creek Foundation, and Brenden-Colson Center foundation to RCS. - Rosalie C. Sears, 2730 SW Moody Ave, Portland, Oregon 97201. Mail code: CL4RS. - Declaration of interests: **Rosalie C. Sears**: Consultant: Novartis Pharmaceutical, Larkspur Biosciences. Scientific Advisory Board: RAPPTA Therapeutics. Sponsored Research Support: Cardiff Oncology, Astra Zeneca Partner of Choice grant award. **Gordon Mills**: SAB/Consultant: Amphista, Astex, AstraZeneca, BlueDot, Chrysallis Biotechnology, Ellipses Pharma, ImmunoMET, Infinity, Ionis, Leapfrog Bio, Lilly, Medacorp, Nanostring, Nuvectis, PDX Pharmaceuticals, Qureator, Roche, Signalchem Lifesciences, Tarveda, Turbine, Zentalis Pharmaceuticals. Stock/Options/Financial: Bluedot, Catena Pharmaceuticals, ImmunoMet, Nuvectis, SignalChem, Tarveda, Turbine, Licensed Technology, HRD assay to Myriad Genetics, DSP patents with Nanostring. Sponsored research: AstraZeneca, Nanostring Center of Excellence, Ionis (Provision of tool compounds. **Lisa M. Coussens** reports consulting services for Cell Signaling Technologies, AbbVie, the Susan G Komen Foundation, and Shasqi, received reagent and/or research support from Cell Signaling Technologies, Syndax Pharmaceuticals, ZelBio Inc., Hibercell Inc., and Acerta Pharma, and has participated in advisory boards for Pharmacyclics, Syndax, Carisma, Verseau, CytomX, Kineta, Hibercell, Cell Signaling Technologies, Alkermes, Zymeworks, Genenta Sciences, Pio Therapeutics Pty Ltd., PDX Pharmaceuticals, the AstraZeneca Partner of Choice Network, the Lustgarten Foundation, and the NIH/NCI-Frederick National Laboratory Advisory Committee. **All the others authors** declare no competing interests.

**Keywords:** MYC, PTEN, triple-negative breast cancer, modeling, Heterogeneity

## Abstract

Triple-negative breast cancer (TNBC) patients have a poor prognosis and few treatment options. Mouse models of TNBC are important for development of new targeted therapies, but few TNBC mouse models exist. Here, we developed a novel TNBC murine model by mimicking two common TNBC mutations with high co-occurrence: amplification of the oncogene MYC and deletion of the tumor suppressor PTEN. This Myc;Ptenfl murine model develops TN mammary tumors that display histological and molecular features commonly found in human TNBC. We performed deep omic analyses on Myc;Ptenfl tumors including machine learning for morphologic features, bulk and single-cell RNA-sequencing, multiplex immunohistochemistry and single-cell phenotyping. Through comparison with human TNBC, we demonstrated that this new genetic mouse model develops mammary tumors with differential survival that closely resemble the inter- and intra-tumoral and microenvironmental heterogeneity of human TNBC; providing a unique pre-clinical tool for assessing the spectrum of patient TNBC biology and drug response.

**Statement of significance:** The development of cancer models that mimic triple-negative breast cancer (TNBC) microenvironment complexities is critical to develop effective drugs and enhance disease understanding. This study addresses a critical need in the field by identifying a murine model that faithfully mimics human TNBC heterogeneity and establishing a foundation for translating preclinical findings into effective human clinical trials.

## Introduction

Breast cancer (BC) is clinically characterized by expression of three receptors: estrogen receptor (ER), progesterone receptor (PR) and human epidermal growth factor receptor 2 (HER2). BC is classified pathologically as hormone receptor (HR) positive (ER+/PR+/-), HER2+, or triple-negative breast cancer (TNBC), lacking expression of all three markers [1, 2]. Expression of these receptors has prognostic value. Although TNBC represents a minority of all breast carcinomas, approximately 10-15%, it is a challenge in clinical practice due to the poor clinical outcomes and greater mortality compared with non-TNBC [2–5]. In addition to the aggressive nature of TNBC, the limited targeted therapy options and lack of sensitivity to endocrine agents contribute to significantly shorter disease-free and overall survival (OS) [3, 5]. Although multiple therapeutic options such as anti-PDL1, Sacituzumab, Enhertu and PARP inhibitors are changing the landscape, TNBC is still currently the worst outcome of breast cancer. With current standard therapies, the median OS for the disease is 10.2 months, with a 5-year survival rate of ~65% for patients with regional tumors and 11% for those with disease spread to distant organs [6, 7].

Around 70% of triple-negative tumors are molecularly subtyped by gene expression as basal-like [1, 8, 9], which is characterized by aggressive phenotypes with higher rates of proliferation, poor differentiation and increased metastatic capability [10]. Previous work analyzing gene-expression profiles of TNBC identified the existence of 6 different TNBC molecular subtypes: basal-like 1 and 2, immunomodulatory, mesenchymal, mesenchymal stem-like and luminal androgen receptor [11, 12]. Another more recent classification for TNBC was suggested by Burstein et al., defining four molecular subtypes: luminal androgen receptor, mesenchymal, basal-like immune-suppressed and basal-like immune-activated [13, 14]. These results emphasize the need for an increased molecular understanding of what drives TNBC heterogeneity and how best to treat the diversity of this disease in the patient population. Accomplishing these goals requires robust laboratory models that capture this heterogeneity to support developing new treatment strategies.

Genetically engineered mouse models (GEMMs) that phenocopy breast cancer provides a powerful platform for testing hypotheses regarding tumor development and progression, interaction with the microenvironment and therapeutic response. Numerous GEMMs have been engineered with inducible, conditional, or constitutively active oncogenes or loss of tumor suppressor genes [15–18]. However, these models are often limited by one or more of the following: 1) being driven by alterations in genes not commonly found to be genetically altered in human breast cancer; 2) not progressing to widespread metastatic disease; and/or 3) forming histological features not commonly found in human breast cancer. As such, current TNBC models reflect only particular aspects of this malignancy and lack the molecular complexity of human TNBC disease.

Amplification of the *MYC* oncogene occurs more frequently in TNBC tumors than other BC subtypes, being found in ~60% of samples, and is associated with worse outcomes [19–21]. In addition to genetic gain, our lab has found that increased phosphorylation of MYC at Ser62 (p-S62-MYC) leads to increases in MYC protein stability and transactivation of target genes in human breast tumors and breast cancer cell lines, including triple-negative [22–24]. Upregulation of MAPK and PI3K pathways, frequently observed in BC, leads to increased p-S62-MYC[25]. Furthermore, there is a positive feedback loop where MYC has been shown to stimulate signaling through the PI3K–AKT pathway via up-regulation of micro-RNAs that down-regulate the tumor suppressor phosphatase gene *PTEN* [26–28]. Consistent with these studies, MYC inactivation was reported to be followed by transcriptional upregulation of *Pten* in mouse MYC-driven osteosarcomas [29]. In addition, disruption in *PTEN* occurs frequently in TNBC, and *PTEN* deficiency is linked with an aggressive subgroup of TNBCs [30]. These *PTEN*-deficient TNBCs display high signaling of MYC, WNT and PI3K pathways and increased drug resistance [30]. These findings raise the hypothesis that MYC gain and PTEN loss may cooperate to drive lethal metastatic disease in human TNBC.

Motivated by this, we developed a genetically engineered mouse model to replicate MYC activation and PTEN loss in human TNBC by combining a *Rosa^LSL-Myc/LSL-Myc^;Blg-Cre* strain with the *Pten^fl/fl^*-conditional knockout mouse model, designated Myc;Ptenfl. We show that combination of MYC deregulation and PTEN loss results in accelerated development of metastatic, heterogeneous triple-negative mammary tumors resembling multiple human TNBC subtypes on gene expression profiles. We performed comprehensive histological, molecular, and transcriptional analyses together with immune composition and localization to show that Myc;Ptenfl mammary tumors recapitulate inter- and intra-tumoral heterogeneity and retain many key features of human TNBC subtypes. Analysis of early-stage tumors demonstrated that even from early time points, intertumoral heterogeneity was observed with multiple histologic subtypes of Myc;Ptenfl TNBC. We found that Myc;Ptenfl tumors effectively recapitulated the different levels of immune activity and various other prognostic characteristics of human TNBC. Together, these results highlight the potential utility of our Myc;Penfl TNBC model for assessing the spectrum of patient triplenegative tumor behavior and drug response.

## Results

### Combination of deregulated MYC and PTEN loss in mammary epithelium drives rapid triple-negative mammary tumors

Copy number aberrations (CNAs) of *MYC* are frequent in BC. Examining the METABRIC dataset [31], which includes over 2,400 primary breast cancer tumors, we found that low-level gain and high-level amplification occurs in 48% of all breast cancers (Supplemental Figure S1A). This includes 60% of ER−/HER2+ breast cancers, in line with our previous observation that our MY C;NeuNT (activated HER2) GEM model closely recapitulates human HER2+ breast cancers [23]. Among 309 ER-/HER2- breast cancers, classified as triple-negative breast cancer (TNBC) from 2500 breast cancer patients [31], 57% had *MYC* gain or amplification CNAs (Figure 1A and Supplemental Figure S1A). These *MYC* CNAs were associated with increased MYC mRNA (Figure 1B) and decreased survival (Supplemental Figure S1B, HR=1.4, 95% CI=1.0-1.9, p<0.05).

**Figure 1:**
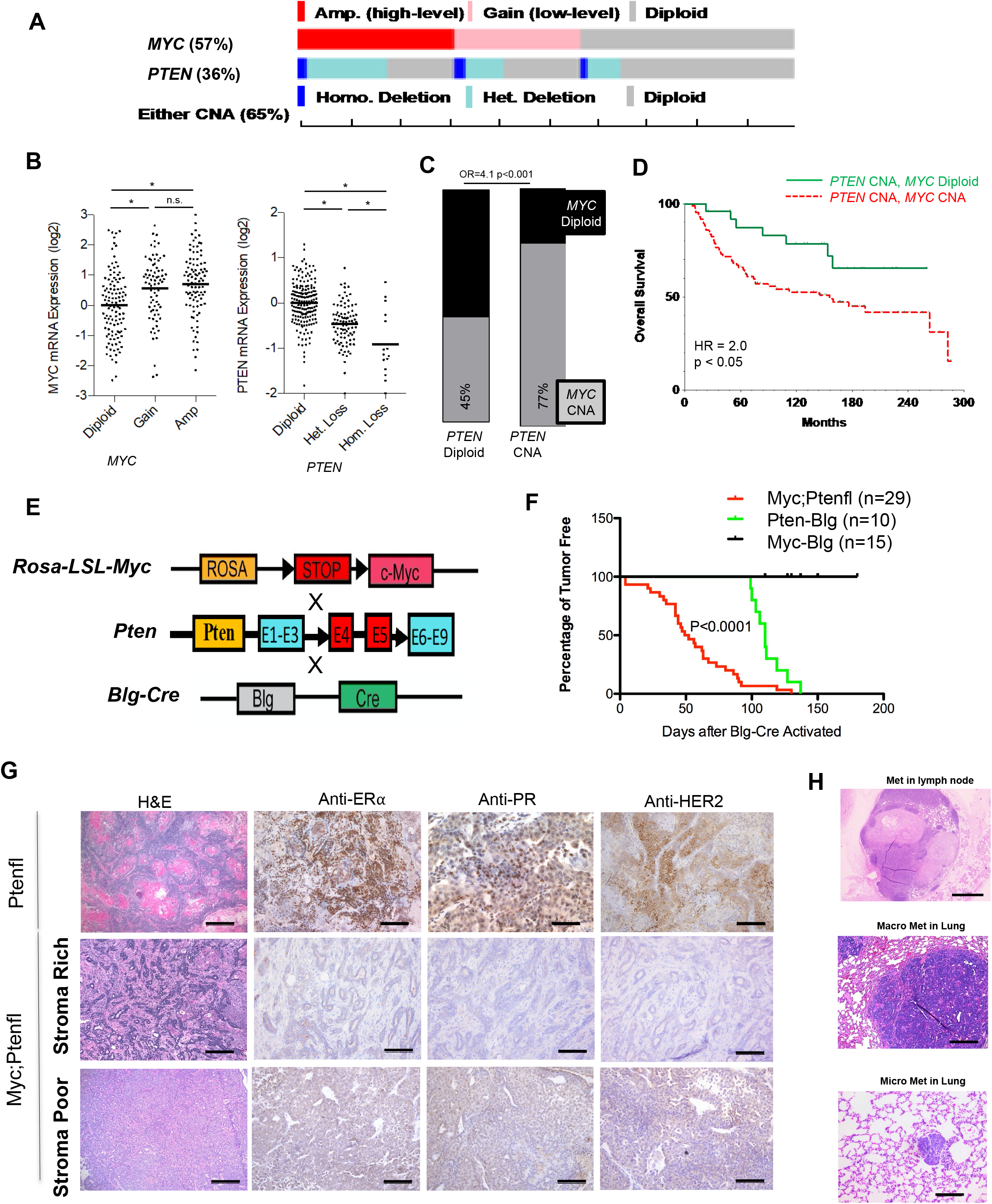
Deregulated *Myc* combination with delated *Pten* in mammary gland accelerate triple-negative mammary tumorigenesis. **A.** and **B** MYC expression and shallow or deep deletion of PTEN in 309 ER- / HER2- patients from 2500 breast cancer patients - METABRIC Data. **C**. Total 101 PTEN deletion in 309 ER- / HER2- patients with 77% MYC amplification or gain; rest 198 ER- / HER2- PTEN expressed patients with 45% MYC amplification or gain. **D.** survival in all 2500 breast cancer patients compare with 12 ER- / HER2- patients(red dotted line) with MYC amplification or gain. **E**. Diagram for generation of Myc;Ptenfl (Rosa-LSL-Myc;Pten^flox/flox^;Blg-Cre) mice breeding the Rosa-LSL-Myc and Pten conditional knockout mice with Blg-Cre transgenic mice. **F**. Mammary gland tumor incidence from Myc (Rosa-LSL-Myc;Blg-Cre), Ptenfl(Pten^flox/flox^;Blg-Cre) and Myc;Ptenfl (Rosa-LSL-Myc; Pten^flox/flox^;Blg-Cre) mice. **G**.H&E staining for Myc;Ptenfl tumors, and Immnuhistochemistry staining with anti-PR, anti-HER2 and Anti-estrogen receptor a (ERa) in 27 mammary gland tumors from Myc;Ptenfl mice and 10 tumors from Ptenfl mice. Scale bar =100μm. **H**. H&E staining for macro lymph node metastisis and micro lung metastisis from 38 Myc;Ptenfl mice and 19 Ptenfl mice. Metastisis in Lymph node scale bar = 1mm, Macro met in Lung scale bar =200 μm, micro lung metastisis scale bar = 100 μm. Pten;Blg 3/19=16%. Myc;Ptenfl SR: Macro 9/23=39%; Micro 3/23=13%. Myc;Ptenfl SP: Macro 5/15=33.3%; Micro 4/15= 26.7%

Breast cancers frequently contain one or more mutations affecting the AKT signaling pathway, including positive regulators, such as *PIK3CA*, and negative regulators, such as the tumor suppressor phosphatase *PTEN*. Among TNBCs, 36% had undergone either loss of heterozygosity (LOH) or homozygous deletion for *PTEN* (Figure 1A), resulting in decreased *PTEN* mRNA expression (Figure 1B). *PTEN* and *MYC* CNAs frequently co-occurred (Figure 1C, Odds Ratio (OR) = 4.1, p<0.001), and 65% of TNBCs had altered *MYC*, altered *PTEN*, or both (Figure 1A). *MYC* CNAs in the presence of *PTEN* loss correlated with poor survival (Figure 1D).

Therefore, we investigated the *in vivo* combination of MYC deregulation and PTEN loss in the mammary gland. We generated Myc;Ptenfl (*Rosa^LSL-Myc/LSL-Myc^;Pten^flox/flox^;Blg-Cre*) mice by crossing the *Rosa^LSL-Myc/LSL-Myc^* mice that express Cre-inducible *Myc* from the *Rosa26* locus, which results in constitutive MYC expression but at physiological levels relevant to human disease [32], with *Pten^flox/flox^* mice for Cre-inducible knock-out of *Pten* [33] (Figure 1E). The *β-lactoglobulin-Cre* (*Blg-Cre*) transgenic mice were used for mammary-specific Cre expression during late pregnancy and lactation [34]. We monitored tumor development in female mice that had passed two cycles of pregnancy/lactation to induce *Blg-Cre* activation at around 10-12 weeks of age. We compared tumor-free survival of the Myc;Ptenfl mice relative to *Myc* deregulated only and *Pten* loss only mice, all in a FVB genetic background. Mice bearing only the deregulated *Myc* did not develop mammary tumors by 24 weeks after *Blg-Cre* activation (Figure 1F), consistent with our previous work using *Rosa^LSL-Myc/LSL-Myc^; WAP- or Blg-Cre* mice where we found low-level expression of wild-type *Myc* from the *Rosa26* locus to be insufficient to drive tumorigenesis [23, 32]. *Pten* loss only mice developed mammary tumors between 110 and 140 days post *Blg-Cre* activation. Combination of deregulated *Myc* with *Pten* loss substantially accelerated tumorigenesis, and these Myc;Ptenfl mice rapidly developed mammary tumors between 4 days and 135 days, average 50 days, post *Blg-Cre* activation (Figure 1F). These synergistic effects were observed with 100% penetrance in FVB background Myc;Ptenfl mice.

To directly examine what subtype of BC these mammary tumors model, we isolated tumors from the *Pten* loss only and Myc;Ptenfl mice and stained for ER, PR, and HER2. We found that *Pten* loss only tumors express ERa, PR, and HER2 receptors and show adenosquamous histology, while combination Myc;Ptenfl mammary tumors are 100% triple-negative for these markers; they do not express ERa, PR, or HER2 (Figure 1G). In addition, and similar to human TNBC, Myc;Ptenfl tumors show histologic heterogeneity with distinct tumor morphology and degrees of stromal involvement not observed with *Pten* loss alone (Figure 1G). The Myc;Ptenfl tumors, based on stromal desmoplasia, fall into two broad classes: stromal rich (SR), which has abundant stroma and displays more heterogenous features including lobular, squamous and metaplastic phenotypes; and stromal poor (SP), which is a solid-pattern invasive ductal carcinoma, the most common architectural pattern seen in TNBC and accounts for 55% of breast cancer morphology at diagnosis [35] (Figure 1G). Preliminary IHC also revealed that SR tumors express higher stromal collagen by Trichrome stain and smooth muscle actin (SMA), basal marker Cytokeratin 5 (KRT5) and phosphorylated Smad3, which can promote epithelial-mesenchymal transition [36], compared with SP tumors. While the model is driven by deregulated *Myc* gene expression, expression of the post-translationally activated S62 phosphorylated MYC was increased in SP relative to SR tumors (Supplemental Figure S1D). pS62-MYC proteins are commonly overexpressed in human breast cancers, and associated with aggressive tumors in mice [23, 24, 32].

We also observed that along with accelerated tumor onset and triple-negative status, Myc;Ptenfl tumors were more metastatic at the IACUC-defined endpoint, with a 52% and 60% metastasis rate to the lymph node and/or lung, in SR and SP, respectively, compared with only 16% overall metastatic rate for Ptenfl endpoint tumors (Figure 1H, and Supplement Figure S1E). Together, these data indicate that the Myc;Ptenfl mice are an *in vivo* model of heterogeneous, aggressive TNBC.

### Myc;Ptenfl tumors show molecular and histologic subtype heterogeneity

It is well understood that TNBC is a heterogeneous disease, with intertumoral heterogeneity observed within the TNBC subtype [37]. We performed gene expression profiling on 13 Myc;Ptenfl tumors, and 3 control (no Cre) normal mammary glands by RNA sequencing (RNA-Seq) to examine the molecular characteristics of this murine model of triple-negative breast cancer. Interestingly, unsupervised hierarchical clustering on normalized gene expression data recapitulated the SP and SR histologic groupings (Figure 2A, clusters 1 vs 2), with several molecular subclusters for SR (cluster2), which is the more heterogenous subtype. These two distinct tumor clusters were also revealed using unsupervised Principal Components (PC) analysis and clustering with the 1200 most variable genes, both of which split the Myc;Ptenfl tumors into two distinct tumor clusters (Supplemental Figure S2A and B). Histopathologic analysis by two different board-certified pathologists, blinded to genotype, recorded several histologically distinct features in the heterogeneous stromal rich tumors, that molecularly subclustered together within the SR cluster (Figure 2A). These included invasive ductal carcinoma (IDC) with lobular features, sufficient squamous differentiation to be classified as IDC with squamous features, and metaplastic IDC with mixed cell phenotypes including spindle-like cells (Figure 2B and C). SP tumors were marked as solid invasive ductal carcinoma with less stromal desmoplasia (Figure 2B and C). In histological analysis of 123 Myc;Ptenfl tumors, we found a histologic distribution of 77% SR subtypes (predominately IDC with lobular features 60%, squamous 15% and metaplastic 2%) and 23% SP subtype tumors (Figure 2C). Thus, they follow human tumors where metaplastic is rare and accounts for 0.2–5% of all breast cancers [38]. Interestingly, mice bearing SR tumors developed tumors earlier than SP, about 45 days after *Blg-Cre* activation (Figure 2D), however these SR tumors took an average of 45 days to reach 2cm diameter tumors (Figure 2E). Whereas mice bearing SP tumors took an average of 90 days after *Blg-Cre* activation to develop tumors (Figure 2D), but these SP tumors grew faster than SR tumors, with an average of 30 days to reach the 2 cm diameter end-point size (Figure 2E). Thus, SP is the more aggressive subtype with the worst overall survival compared with SR (Figure 2E).

**Figure 2:**
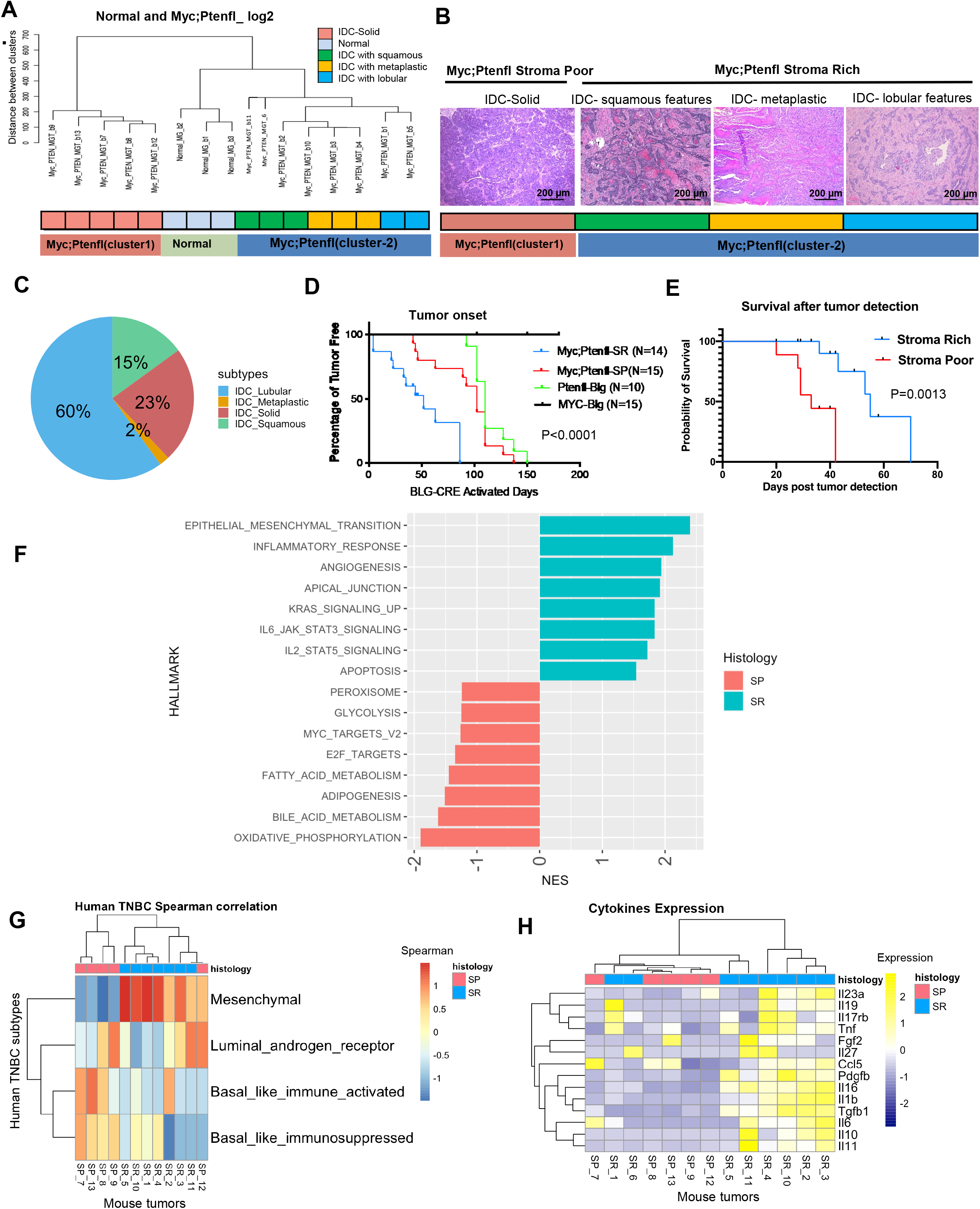
Multiple molecular and Histologic subtypes are present in Myc;Ptenfl tumor with human TNBC subtype-specific transcriptomic signatures. **A.** Unsupervised hierarchical clustering of RNA expression from 13 Myc;Ptenfl mammary gland tumors and 3 controls (normal mice mammary gland). **B**. On the left; histology of Myc;Ptenfl cluster-1 termed as stromal poor (SP) group, on the right; multiple histological subtypes of Myc;Ptenfl cluster-2 termed as stromal rich (SR) group; squamous differentiation (IDC with squamous features), or metaplastic IDC or lobular cells (IDC with lobular features). **C**. Pie chart of an extended histological analysis of 123 tumors from Myc;Ptenfl mice showed the frequency of histological subtypes. **D.** Mammary gland tumor incidence from Myc;Ptenfl stromal rich and stromal poor groups. **E.** Survival after tumor detection for stromal rich tumor bearing mice verse the mice with stromal poor tumor. **F.** Gene set enrichment analysis showing the top 8 upregulated (green) and top 8 downregulated hallmarks in SR compared with SP (orange) visualized as a par plot. X-axis is normalized enrichment score (NES) and y-axis is hallmark gene set. **G.** Applying 77 gene signature from human TNBC to gene expression levels of Myc;Ptenfl mouse samples in correlation analysis to define four prognostically-distinct human TNBC subtypes; basal-like immune-activated (BLIA), basal-like immunosuppressed (BLIS), luminal androgen receptor (LAR), and mesenchymal (MES). **H.** Cytokines expression from the RNA expression data of Myc;Ptenfl tumors in stromal rich compare to stromal poor tumors.

We investigated whether these histologically heterogenous Myc-Ptenfl subtypes originated from similar or distinct tumor phenotypes at an early stage. For this, we examined the phenotypes, and tumor histology of 32 small (dimeter <3mm) tumors. This analysis revealed that even at an early stage, heterogeneous histologic subtypes were observed including solid ductal carcinoma and SR histologies including papillary, lobular and adenosquamous, with more stromal desmoplasia and high expression of the basal marker KRT14, versus solid tumors with lower expression of the basal marker KRT14 (Supplemental Figure S3A and B). Also, Myc;Ptenfl mice even at an early stage, develop triple-negative mammary tumors; they are negative for ERa, PR and HER2 expression (Supplemental Figure S3C). These data suggest that this model may provide a unique resource for examining early events that generate distinct triple-negative breast cancer subtypes.

We next used Genes Set Enrichment Analysis (GSEA) and the Hallmarks of Cancer database to examine molecular pathways active in the Myc;Ptenfl SR vs SP tumors. This demonstrated that the SR tumors are enriched for pathways related to epithelial mesenchymal transition (EMT), angiogenesis, KRAS, IL2 _STAT5 signaling, apical junction, inflammatory response and apoptosis (Figure 2F and Supplemental Figure S2C, supplemental dataset 1), whereas the SP tumors are enriched in gene sets related to metabolism, glycolysis, fatty acid metabolism and oxidative phosphorylation (OXPHOS) (Figure 2F and Supplemental Figure S2C, supplemental dataset 1). OXPHOS has been shown to be important for the production of biosynthetic intermediates necessary to support the rapid proliferation of cancer cells and associated with high lethality in TNBC [39], which could explain the high growth rate and poor prognoses seen in the Myc;Ptenfl SP subtype. We also examined the expression of genes related to different differentiation states that are enriched in luminal, basal or mesenchymal differentiated breast cancer cell lines [40, 41]. Consistent with IHC expression, SR tumors had higher levels of mesenchymal and basal gene expression, with lower luminal gene expression (Supplemental Figure S2D). Whereas SP tumors had higher luminal gene expression. Lehmann and colleagues identified a human subtype of TNBC characterized by expression of luminal genes termed the LAR subtype. This luminal gene expression-based TNBC subtype was subsequently confirmed by other groups [11, 13].

We examined whether the intertumoral heterogeneity observed in our TNBC mouse model is representative of described molecular subtypes of human TNBC. We used the recent human TNBC subtyping method, developed by Ding et al, which uses unique 77-gene centroid signatures to differentiate between four human TNBC subtypes [14]. Consistent with the differentiation states analysis, the correlation between our Myc;Ptenfl tumor samples and the centroids demonstrated that the SR tumors mostly correlated with the human TNBC Mesenchymal (MES) subtype, whereas SP tumors were mixed basal and luminal and mostly correlated with basal-like immune suppressed and/or luminal androgen receptor (LAR) human TNBC (Figure 2G, and supplemental dataset 1). However, IHC staining for AR indicated <1% AR staining in both SP and SR tumors (Supplemental Figure S2E).

GSEA results (Figure 2F) indicated that Inflammatory Response is one of the top Hallmarks of Cancer enriched in the SR phenotype, which has a better survival post tumor detection. Consistent with this, high levels of infiltrated inflammatory immune cells, in particular lymphocytes, has been shown to be associated with improved overall survival in human TNBC [42]. To investigate if core immune signals are differentially expressed across Myc;Ptenfl tumor subtypes, we evaluated the inflammatory cytokines expression among the subgroups of Myc;Ptenfl mouse model tumors. The SR subtype had markedly higher cytokine gene expression than the SP subgroup (Figure 2H).

### Multiplexed immunohistochemistry platform identifies distinct immune complexity profiles across Myc;Ptefl subtype tumors

Considering the important role of the tumor immune microenvironment in prognosis, we utilize a multiplexed immunohistochemistry (mIHC) approach to characterize the immune contexture and the spatial distribution of immune cells among the subgroups of the Myc;Ptenfl mouse tumors. The mIHC platform comprises a validated panel of 23 antibodies in a sequential staining method for identification of lymphoid and myeloid immune cell lineages, epithelial markers, and functional markers in a single FFPE tissue section (Figure 3A and B, and supplemental dataset 2) [43, 44]. Qualitatively, SR tumors show higher infiltration of lymphoid and myeloid immune cell lineage, including T-cells markers (CD3, CD8 and CD4), B cells marker (CD45R), Treg cell marker (Foxp3), and macrophages marker (CD68) than SP tumors (Figure 3A and B).

**Figure 3:**
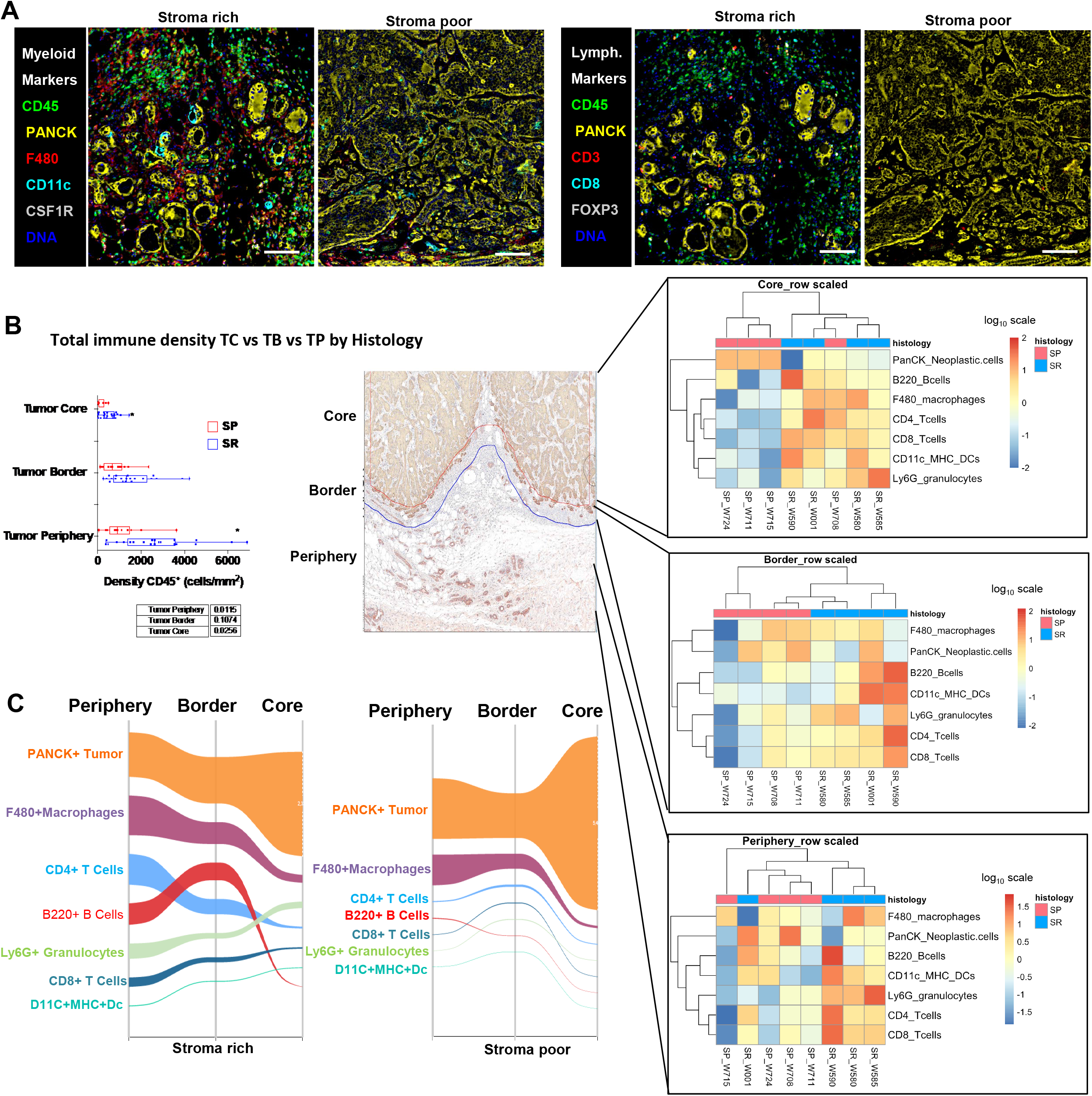
multiplexed immunohistochemistry platform defined immune contexture across Myc;Ptenfl subtypes. **A.** Example of mIHC images from Myc;Ptenfl stromal rich tumors and stromal poor tumors with Myeloid markers expression on the left and Lymphoid markers expression on the right. Scale bars = 100 μm. **B.** Total immune cells by CD45^+^ density in stromal rich tumor periphery, border and core compare to stromal poor tumor in Myc;Ptenfl mice. Tumor border was determined by CD45 and PANCK expression (supplemental S4 A and B). On the right; Heatmap of raw density values displayed on log_10_ scale in each periphery, border and core of Myc;Ptenfl tumor subtype. **C.** Sankey diagrams showing the distribution of each immune lineage population in stromal rich (left) and stroma poor (right) tumors; periphery, border and core.

Quantitative assessment and spatial, *in situ* localization are vital for imaging-based biomarker exploration of the tumor microenvironment; thus, we utilized an image analysis pipeline [45], for evaluation of the 23-plex images. Additionally, because immune cells are often excluded from the tumor core and sequestered at tumor borders or beyond [46], we elected to frame our analysis around three spatial categories: tumor periphery, tumor border, and tumor core (Figure 3B, supplemental Figure S4). As predicted, the lowest density of total CD45+ immune cells was found in the tumor core, for both subgroups, and an increasing density of immune cells was observed in the border and periphery (Figure 3B). However, SR tumors had significantly higher total immune cell density in all tumor core, border and periphery compartments compared to SP. Next, we employed an unsupervised hierarchical clustering approach, and observed two distinct immune complexity profiles for SR and SP tumors across spatial compartments, where lymphoid and myeloid lineage cells were differentially present between the groups, indicating distinct immune contexture by subtype (Figure 3B). When performing supervised analyses, we observed that SR tumors had trending higher densities of Ly6G^+^ granulocytes, B220^+^ B cells, CD11c^+^ DCs, and CD4^+^and CD8^+^ T cells in all spatial compartments, with significantly higher density of Ly6G^+^ granulocytes and CD8^+^ T and DC cells in tumor border and periphery, respectively (Supplementary Figure S4C-E). The proportion of FoxP3^+^ Tregs within the CD4^+^ T cells was greater in SR (Supplementary Figure S4F and G). Sankey diagrams showing relative density of the indicated cell types by spatial compartment indicate that PanCK^+^ neoplastic cells dominate in all spatial categories with an increase toward the tumor core in both subtypes (Figure 3C). As expected, the SP group shows very high abundance of PanCK^+^ neoplastic cells in the tumor core. Notably, CD4^+^ and CD8^+^ T cells (light blue and dark blue lines, respectively) are most abundant in the periphery and border, particularly in SR tumors, dropping in abundance dramatically in the tumor core, while macrophages dominate in all spatial categories (maroon line) (Figure 3C). Clearly, SR tumors contain higher abundance of multiple immune cell types and trends towards more lymphocytes compared to SP tumors. Particularly, CD8^+^ T cells are significantly higher in the SR tumor periphery than SP periphery (Supplementary 4E, p-value= 0.05). Indeed, several studies have shown that high levels of stromal T cells are associated with improved overall survival in human TNBC [13, 42, 47]. These findings indicate that the Myc;Ptenfl model may provide a unique tool for assessing a spectrum of immune cell phenotypes in triple-negative tumor behavior and drug response.

### Morphological feature extraction identifies shared morphologies between human breast cancer and Myc;Ptenfl mammary tumors

To compare the morphological features of Myc;Ptenfl TNBC mouse with human patient TNBC, we compared several tissue microarrays (TMAs), one generated from our TNBC mouse model and two generated from human breast cancer cases from Vanderbilt University Medical Center. The human TMAs are comprised of 60%TNBC, 30% ER+ and 10 %HER2+ disease. To assess the morphological features of each tumor core, H&E core images from the TMAs were tiled into 28 regions per core. We used a variational autoencoder (VAE) [48, 49] - an unsupervised deep learning (DL)-based method for representation learning and feature extraction (Figure 4A, UMAP embedded image shown in Supplemental Figure S5A). From each normalized H&E tile, we extracted a morphological feature vector to establish comparison between each tile, cores, and tissue origins. Tiles were compared using UMAP embedding [50, 51] as well as k-means clustering analysis of the latent feature vectors. The regional H&E tile images on UMAP showed distinct morphological feature differences, and density maps for human and mouse tiles showed overlapping regions in UMAP embedding space, highlighting shared morphological features between human and mouse TNBC (Figure 4B). K-means clustering identified eight representative groups (Figure 3C). Representative tile images near the cluster center and single high-resolution tiles within each cluster are shown to illustrate the distinct morphologies of each cluster. Each cluster showed various morphology such as carcinoma with discohesive growth pattern (cluster a), carcinoma with thin fibrotic stroma and hyperchromatic nuclei (cluster b), coagulative necrosis (cluster c), IDC high grade (cluster d), stromal inflammatory cell infiltration (cluster e), fibrotic stroma (cluster f), sarcomatoid change (cluster g), tumor with hyperchromatic and course chromatin with frequent atypical mitoses (cluster h) (Figure 3D). Also, the relative abundance of human and mouse TMA tiles was calculated for each cluster to show the cluster’s tissue of origin and makeup (Figure 3E). Overall, quantitative cluster analysis shows a high level of overlap between human and mouse tiles. The most prominent human morphologies (clusters a, d, and f), which comprise 72% of all human tiles, also have a high representation of mouse tiles. Clusters that show a high-class imbalance (clusters b and e), comprise a much smaller portion of the total tile population. Because human TMAs included both TNBC and non-TNBC, we ran the analysis on human TMAs, comparing the VAE results based on the BC subtype. We found that ER+ and TNBC subtypes highly overlaped with each other, indicating that the different architectural level features identified by the VAE are not specific to a single subtype (Supplemental Figure S5B).

**Figure 4:**
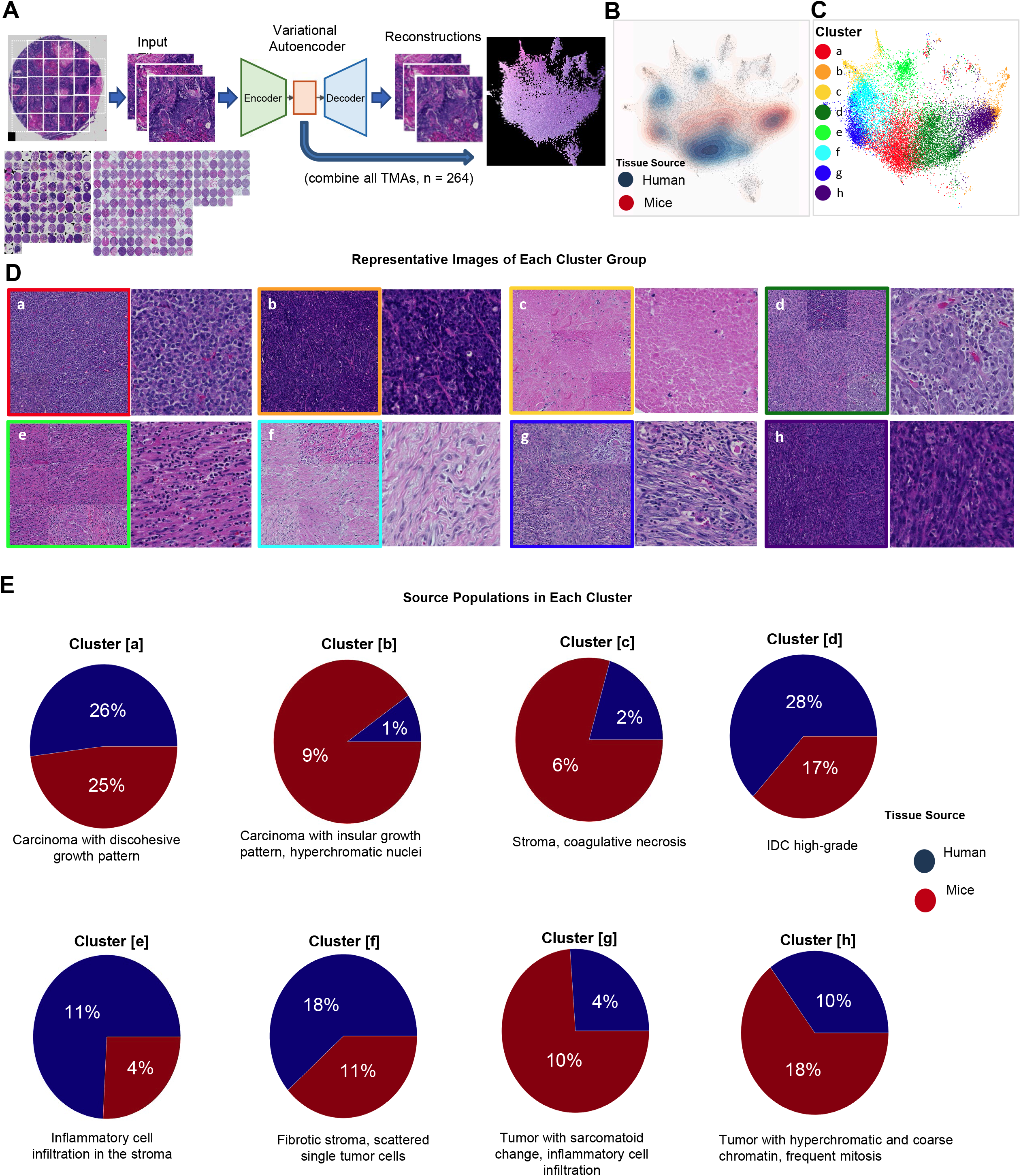
Analysis of Shared Morphologies Between Human breast cancer and Mice TMAs. **A**. Pipeline for generating morphological feature representation using a variational autoencoder (VAE). Tiles from both mice and human TMAs are used to train a VAE, then a latent encoding vector is computed for each tile. Tiles are compared using UMAP embedding and k-means clustering analysis of the latent features. **B**. Density functions for all human and mice tiles are calculated within the 2-dimensional UMAP space to visually compare overlap in embedding space (corresponding histopathological feature image is shown in Supp Fig 5). **C.** K-means clusters (n=8) are computed using latent features and projected into UMAP space for visualization. Clusters consisting primarily of edge artifacts were excluded from analysis. **D.** For every cluster, the 9 tiles closest to the cluster center (left) and single high-resolution tile image within each cluster (right) are shown to illustrate each clusters dominate morphology; main histologic features of each center:: [a] carcinoma with discohesive growth pattern, [b] carcinoma with thin fibrotic septa and hyperchromatic nuclei, [c] stroma or coagulative necrosis, [d] IDC with high-grade nuclear feature, [e] inflammatory cell infiltration in the stroma, [f] fibrotic stroma, scattered single tumor cells [g] sarcomatoid change of tumor cells and inflammatory cell infiltration, [h] tumor with hyperchromatic and coarse chromatin with frequent atypical mitosis. **E.** Relative abundance of human and mice TMAs are calculated for each cluster using the ratio of tiles in a cluster to total tiles from the given TMA source.

To compare histologic features of mice and human TMAs, a pathologist (E.N.K) manually annotated detailed histologic features of the original mouse TNBC TMA digitized slide (80 TMA cores) (Supplemental Figure S6 and Supplementary dataset 3). The Myc;Ptenfl mice tumors show a diverse histologic spectrum from ductal carcinoma not otherwise specified (NOS) to various metaplastic elements, similar to the typical characteristics of human TNBC [52, 53]. Myc:Ptenfl TNBC tumors showed IDC, histologic grade 2 (low grade nuclear features) (21.25%, 17/80), histologic grade 3 (high grade nuclear features) with marked nuclear pleomorphism, and prominent nucleoli (17.5%, 14/80), solid sheet-like growth pattern without tubule formation (12.5%, 10/80), stromal proliferation with lymphocytic infiltrate (33.75%, 27/80), geographic necrosis (21.25%, 17/80) [54], and a myoepithelial phenotype, which belongs to basal cells, was prominently observed in mouse TNBC (41.25%, 33/80). Like human TNBC, various metaplastic changes such as sarcomatoid change with spindle cells (12.5%, 10/80) and squamoid change with keratin pearls (47.5%, 38/80) were observed [55]. Although it was not identified in the human TNBC TMA cores of this study, clear cell change (11.25%, 9/80)[56] and thick trabeculae suggestive of neuroendocrine differentiation (1.25%, 1/ 80) [57], which are reported as very rare human TNBC, were also observed in the mice TNBC.

Notably, the most abundant histological features in the Myc;Ptenfl SR subtype are: squamoid metaplasia, myoepitheial proliferation and stromal lymphocytic infiltrate with 59.37%, 51.56% and 39% frequency, respectively. Although, these metaplastic tumors characterized by squamous metaplasia have been shown to be highly chemoresistant and aggressive, the stromal lymphocytic infiltrate phenotype has been associated with improved survival [58–62]. In contrast, the most frequent histological features in the Myc;Ptenfl stromal poor phenotype, which has a poorer prognosis, are low and high grade nuclear features and a solid growth pattern with 50%, 37.5%, and 50% frequency, respectively. Indeed, human TNBC studies showed that a solid growth pattern and high grade nuclear features are typically associated with poor TNBC prognosis, resistance to systemic cytotoxic therapy and histological features of aggressive TNBC [58–61]. These results indicate that our mouse model recapitulates the histological heterogeneity and corresponding spectrum of prognostic features seen in TNBC patients.

### Shared Tumor and Microenvironment Cellular Phenotypes in Myc;Ptenfl TNBC Tumors and Human TNBC

Next, we used cyclic immunofluorescence (CyCIF, 20 cell type and phenotype markers were analyzed, see Supplemental dataset 4) to examine tumor and microenvironment cell phenotypes in our TNBC mouse model. CyCIF staining of the Myc;Ptenfl TMA confirmed the generalized histologic SR and SP subtypes with increased stromal cells in the previously designated SR tumors (Figure 5A). Image processing, single-cell segmentation and feature extraction was performed with the mplexable software [63]. Gating on cell-type specific markers determined cell frequencies of epithelial, immune and stromal cells in each tissue (Supplementary Figures S7A-C), and showed SP as more enriched with epithelial cells compared with SR (Figure 5B). In order to compare our data to human TNBC, we obtained a publicly available human TNBC multiplex ion beam imaging (MIBI) dataset [64]. Gating on cell-type specific markers identified epithelial, immune and stromal cells in the human tissue (Supplemental Figure S7D-F). We compared cell frequencies of epithelial, stromal and immune cells between our TNBC mouse model and the human TNBC MIBI data. Clustering of mice and human tissues based on cell frequency and further annotation by the levels of epithelial, immune and stromal non-immune cell types in clusters revealed three general phenotypes: stroma poor (SP), stroma-rich-immune-rich (SR-IR) and stroma-rich-immune-poor (SR-IP) (Figure 5C-E, and Supplementary Figures S8). Further splitting the SP phenotype into SP+ with the lowest stroma demonstrated significant association with the original mouse histology SP and SR subgroup designations (Supplementary Figures S9A-D), in which SP+ samples by CyCIF cell types were also SP by original histology (7 out of 10) and SR histology generally subdivided into SR-immune rich and -immune poor subtypes by CyCIF (Supplemental Figure S9). Furthermore, the mean frequency of epithelial, immune and stromal cell types in each of the three SR-IR, SR-IP, and SP subtypes was similar between mouse and human TNBC (Figure 5E). Interestingly, the SP and SR subtypes were prognostic in human MIBI data, with stroma poor having the worst overall survival compared with the stroma rich subtypes (Figure 5F, Supplemental Figure S8G), consistent with subtype survival in our TNBC mouse model (Figure 2E).

**Figure 5:**
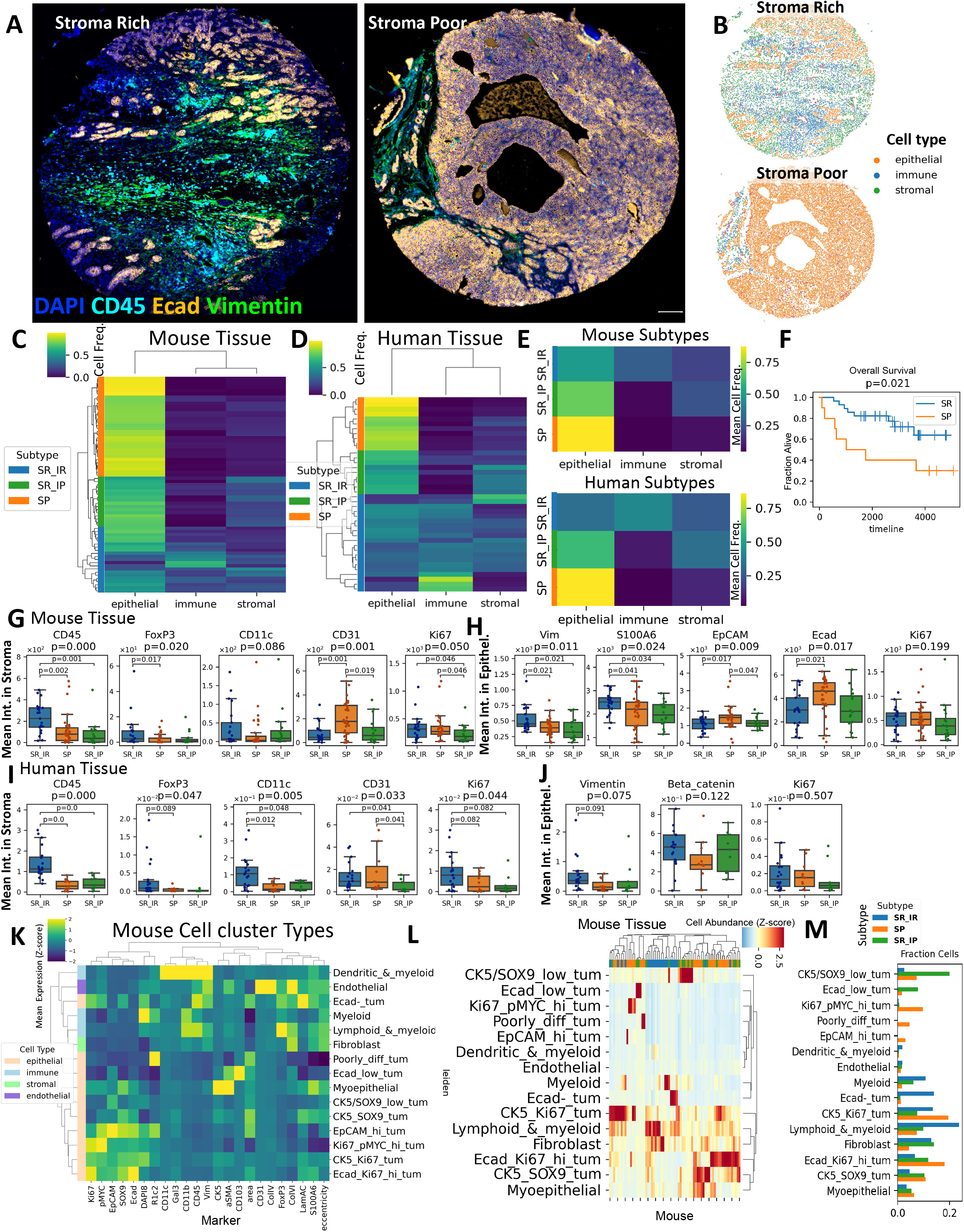
Tumor Microenvironment Subtypes Defined by Multiplex Imaging in Mouse and Human TNBC. **A.** Cyclic immunofluorescence staining of representative tissue microarray (TMA) cores of stroma rich (left) and stroma poor (right) histology subtypes. Scale bar = 130 μm. **B.** Cell type calling defined by gating on cores in (A). **C.** Hierarchical clustering of mouse samples based on cell type frequency in each tumor. **D.** Hierarchical clustering of human samples based on cell type frequency in each region of interest in TNBC tumors imaged with multiplex ion-beam imaging (MIBI). **E.** Mean frequency of cell types in cell frequency-based subtypes in mouse (top) and human (bottom) samples. **C-E.** Heat map row colors: stroma-poor (SP, orange), stroma-rich, immune-rich (SR_IR, blue) and stroma-rich-immune-poor (SR_IP, green) subtypes. **F.** Kaplan-Meier curves of overall survival in cell frequency-based subtypes in human TNBC. Log-rank p=0.021. **G.** Stromal expression of pan-immune (CD45), Treg (FoxP3), dendritic (CD11c), endothelial (CD31) and proliferation (Ki67) markers in mouse subtypes. **H.** Epithelial expression of mesenchymal (Vim, S100A6), epithelial (EpCAM, Ecad) and proliferation markers in mouse subtypes. **I.** Stromal expression of pan-immune, Treg, dendritic, endothelial and proliferation markers in human subtypes. **J.** Epithelial expression of mesenchymal (Vimentin) and proliferation markers in human subtypes. **G-J**. P-values determined by Kruskal–Wallis H test. **K.** Mean marker intensity in each annotated cell type defined by unsupervised Leiden clustering in mouse tissue. **L.** Hierarchical clustering of mouse samples based on detailed cell types. Heat map column colors: stroma-poor (SP, orange), stroma-rich, immune-rich (SR_IR, blue) and stroma-rich-immune-poor (SR_IP, green). **M.** Bar plot of frequency of each cell type in mouse subtypes.

We compared mean expression of key markers in stromal cells and epithelial cells in the three subtypes in human and mouse tumors and found similar subtype-significant differences in compartment-specific marker expression between human and mouse (Figure 5G-J, Supplementary 10A and B). In both human and mouse tumors, the stroma-rich-immune rich subtype had greater stromal pan-immune CD45, T regulatory FoxP3 and dendritic CD11c, consistent with the multiplex IHC platform (Figure 3 and supplementary 4). In mouse SP tumors, the sparse stromal cells expressed relatively more of the endothelial marker CD31 (Figure 5G). Human SP tumors also showed higher stromal CD31 expression (Figure 5H). Stromal proliferation as measured by Ki67 expression differed among the three subtypes in mouse and human tumors, but there was no difference in epithelial Ki67 (Figure 5G-J). Human and mouse epithelial cells from SR-IR tumors showed higher expression of the mesenchymal marker vimentin, higher nuclear eccentricity and lower expression of the epithelial markers E-cadherin and EpCAM compared to SP (Figure 5H, supplemental Figure S10A), consistent with the mouse bulk RNAseq data demonstrating that SR tumors had more mesenchymal gene expression and subtyped as human TNBC mesenchymal (Figure 2G and Supplemental S2D). S100A6 was elevated in the SR-IR mouse epithelial cells, while β-catenin was elevated in the SR-IR human epithelial cells (Figure 5H, J). Since previous work showed S100A6 promoting EMT through β-Catenin in pancreatic cancer cell lines [65], the S100A6-high/β-Catenin-high phenotype in mouse/human SR-IR tumors may reflect a shared mechanism of EMT. Similar to the original IHC staining (Supplemental Figure S1), the histologically designated SR subtype showed high expression of αSMA and the basal marker CK5 compared to the SP, while Phospho-MYC was higher in the SP subtype epithelial cells (Supplemental Figure S10C), consistent with IHC results (Supplemental Figure S1D).

To take advantage of the power of the multiplex imaging data to study co-expression of phenotypic markers within their tumor context, we performed unsupervised clustering of single cells with the Leiden algorithm based on 20 markers plus nuclear area and nuclear eccentricity in the mouse TMA. This resulted in 21 clusters (Supplemental Figure 7C) which, after artifact removal and combining of similar clusters, produced 14 distinct annotated cell type clusters (Figure 5K). These cell types captured intratumoral heterogeneity observed in the images (supplemental S11A and B). Mouse TMA tissues were then hierarchically clustered based on the z-scored cell abundance of these cell types, revealing phenotypic heterogeneity within the stroma-poor and stroma-rich immune-poor subtypes (Figure 5L). Conversely, most of the stroma-rich, immune-rich tissues clustered together, driven by high abundance of fibroblasts and immune cells in the tissues (Figure 5L). Consistent with our other analyses, SR-IR tumors were enriched in immune cells, had fewer Ki67+ proliferating tumor cells and also contained E-cadherin negative epithelial cells, fitting with a mesenchymal phenotype, while SP tumors were enriched in Ki67+tumor cells and de-enriched for E-cadherin low/negative epithelial cells and fibroblasts, consistent with their fast-growing, solid pattern (Figure 5M). Single cells from the human MIBI data were clustered with the Leiden algorithm resulting in 25 cell type clusters that were annotated as epithelial, immune, or stromal (Supplemental Figure S7F). Similar to mouse, human SR-IR tumors were enriched in immune cell type clusters (Supplemental Figure S11C-E). Similar to mouse tissues, hierarchical clustering of human samples based on these cell types revealed heterogeneity within the stroma-poor and stroma-rich immune poor subtype while SR-IR tissues clustered together (Supplemental Figure S11D).

### scRNA-seq reveals tumor subtype-specific distributions of cell states

To better understand the cellular heterogeneity within Myc:Ptenfl tumors we performed deep single cell RNA sequencing (scRNA-seq) in which 11 tumors from 6 mice were hashtag oligonucleotide (HTO) multiplexed into sequencing libraries (4 SP, 7 SR assigned by pathologist from H&E) (Supplemental Figure S12). Ambient mRNA contamination was corrected with the R package SoupX [66]. After HTO demultiplexing, in silico doublet identification and quality control thresholding we retained a total of 14,042 cells, ranging from 490 to 5995 cells per tumor, with a mean of 1,198 unique genes and 3,394 Unique Molecular Identifiers (UMI) per cell (Methods, Supplemental figure S13). Simultaneous linear dimensionality reduction and integration across libraries to remove technical artifacts related to library preparation was performed with the rliger implementation of iNMF (Methods, Supplemental figure S14 A and B) [67].

Clusters of cells were identified using the Leiden algorithm applied to iNMF embeddings with parameters optimized for maximum silhouette width [68] (Bluster https://doi.org/doi:10.18129/B9.bioc.bluster). Each cluster was then manually assigned a cell lineage (epithelial, lymphoid, myeloid, fibroblast, endothelial, and perivascular) based on aggregated expression of canonical markers (Figure 6A, supplementary dataset 5). We found that a single cluster (cluster 14) was defined by expression of cell cycle related genes rather than lineage signatures, and this cluster was further subclustered and each subcluster was assigned to an appropriate lineage using the same schema as the primary clusters (Supplemental figure S14 C-E). Investigation into the relative proportion of lineage representation in histologically defined stromal poor and stromal rich tumors showed, consistent with CyCIF, that SR tumors subdivided into tumors with high representation of both immune and non-immune stromal cell types (Stromal-Rich Immune-Rich, SR-IR) and tumors which had high representation of non-immune stromal cells but low representation of immune cells (Stromal-Rich Immune-Poor, SR-IP) (Figure 6 B).

**Figure 6:**
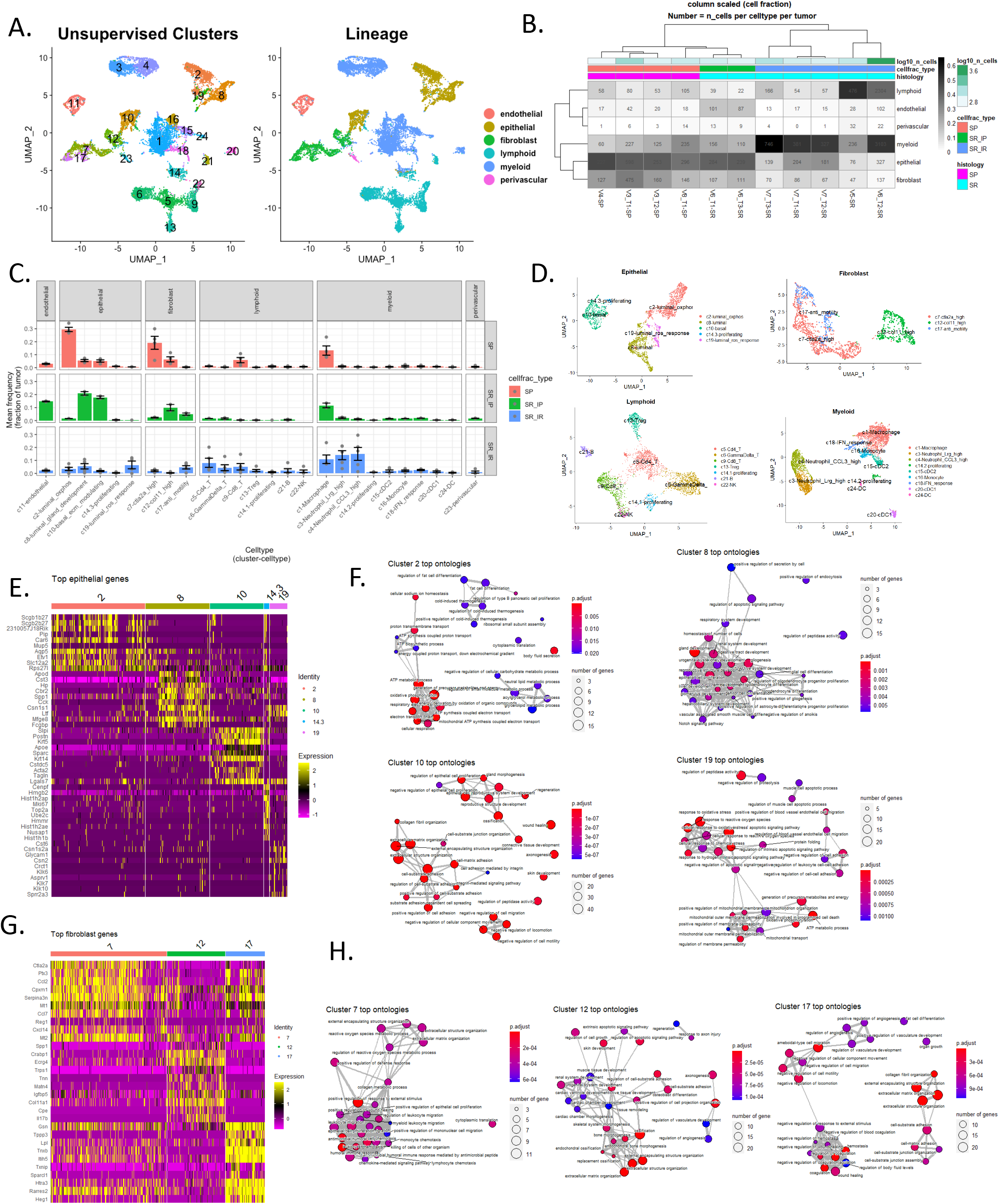
MYC-PTENfl subgroups showed distinct cell type and cluster genes enrichment with similar architecture to human TNBC. **A.** UMAP showing scRNA-seq data from Myc;Ptenfl model. Color coded by either lineage or unsupervised cluster. UMAP was computed on 50 iNMF factors. Lineage was manually assigned using a canonical celltype markers (supplemental dataset table 5) for unsupervised clusters. Unsupervised clusters were identified using the Leiden algorithm to the same 50 iNMF factors (Resolution = 0.45, optimized for silhouette width, supplemental figure 14). **B.** Distribution of cells found in each tumor. Heatmap color represents the proportion (fraction of total cells recovered) and inset number is the number of cells. Histology represents the two classes assigned by a pathologist (SP: Stromal Poor, SR: Stromal Rich). For Cell fraction type the stromal rich subtype was further divided into SR-IR: Stromal Rich Immune Rich, and SR-IP: Stromal Rich Immune Poor). **C.** Relative abundance of clusters for each Myc;Ptenfl tumor subtype. Mean frequency is the arithmetic mean across all tumors within that subtype, error bars represent SEM. **D.** UMAP visualizations of individual lineages. UMAPs and clustering were computed using the same 50 factors as global analysis (Figure 6 A, Supplemental Figure 14 C). **E.** Heatmap showing the top 10 uniquely upregulated genes for each epithelial cluster (min.pct = 0.1, avg_log2FC > 0.5, Bonferroni corrected p <= 0.05). **F.** Enrichment maps showing the top 30 enriched ontologies for each epithelial cluster visualized as a network. Size of point indicates number of genes within the ontology that were uniquely upregulated in that cluster. Edges connect any ontologies with a Jaccard similarity greater than 0.2, and edge width scaled to Jaccard similarity. **G.** Heatmap showing the top 10 uniquely upregulated genes for each fibroblast cluster (min.pct = 0.1, avg_log2FC > 0.5, Bonferroni corrected p <= 0.05). **H.** Enrichment maps showing the top 30 enriched ontologies for each fibroblast cluster visualized as a network. Size of point indicates number of genes within the ontology that were uniquely upregulated in that cluster. Edges connect any ontologies with a Jaccard similarity greater than 0.2, and edge width scaled to Jaccard similarity.

Within each cell lineage we performed differential expression analysis to identify uniquely upregulated genes for a given cluster, separating them from other same-lineage clusters. Putative cell types were assigned using the SCSA approach and then manually refined using a literature mined set of cell type associated genes (Figure 6C and D, supplemental figure S15 A-C, supplementary dataset 5)[69, 70]. Enrichment of Gene Ontology analysis was performed on the differentially upregulated genes for each cluster (average log2 Fold Change > 0.5 and Bonferroni adjust p <= 0.05) and these clusters were assigned names to reflect the themes of their most enriched ontologies (Figure 6C and D). When we compared the proportions of each cell type within a lineage across the three phenotypes, we observed clear distinctions in epithelial and fibroblast cell type proportions and we hypothesized that the different epithelial and fibroblast states may be involved in either immune exclusion or recruitment (Figure 6C and D, supplemental figure S15D). We found that the epithelial cluster 2 was the dominant epithelial state for SP tumors (71% of SP epithelial cells) and this cluster was enriched for programs related to altered metabolism including upregulation of oxidative phosphorylation related ontologies (Figure 6E and F, supplemental figure S15D) and luminal markers (supplemental Figure S15E), which provides further evidence for SP tumors recapitulating OXPHOS high luminal human TNBC subtypes. SR-IP tumors instead were mostly comprised of epithelial cells belonging to either cluster 8 or 10 (51% and 44% of SR-IP epithelial cells respectively, Figure 6C and D, supplemental figure S15D). Epithelial cluster 8 was enriched for ontologies related to gland development and had high expression of several luminal genes (supplemental figure S15E) [11]. Epithelial cluster 10 had enriched ontologies associated with extracellular matrix modulation and high expression of Krt5 and Krt14, both of which are associated with a basal-like TNBC subtype (Figure 6E and F, supplemental figure S15E) [11]. Hence, SR-IP tumor are mixed basal and luminal TNBC. The final epithelial cluster 19 was found at the highest rate in SR-IR tumors (27% of SR-IR epithelial cells, <1% of SR-IP epithelial cells, 2% of SP epithelial cells) and had enriched ontologies for reactive oxygen species response and regulation of apoptotic signaling pathways (Figure 6C-F, supplemental figure S15D).

The Myc;Ptenfl tumors showed a similar subtype-specific enrichment of specific fibroblast cell type clusters. SP tumors were enriched for cluster 7 fibroblasts, which had high Ctla2a expression (Figure 6C, G and H). Ctla2a high in this cluster has been shown to be associated with immune suppression by induction of apoptosis for T-cell lymphocytes [71]. SR-IP tumors were found to have a higher fraction of cluster 12 fibroblasts which uniquely expressed the CAF marker Col11 and had upregulated gene activity related to ossification and apoptosis pathway regulation (Figure 6C, G and H) [72]. The final fibroblast cluster 17 was found at a higher rate in both the SR-IP and SR-IR subtypes, and this cluster was transcriptionally similar to cluster 7 but with the added signaling related to ameboid cell migration and coagulation (Figure 6C, G and H).

Consistent with mIHC, macrophage cells were the most abundant immune cells across Myc;Ptenfl subtypes. The SR-IR subtype had a higher fraction of cells coming from both lymphoid and myeloid lineages when compared to SR-IP and SP tumors. The SR-IR tumors also had a much higher recovery of neutrophils in clusters 3 (Lrg high) and 4 (Ccl3 high). Lrg expression increases during neutrophil differentiation and extracellular release of Lrg from neutrophils was found to be pro-angiogenic [73, 74]. Ccl3 is considered a pro-tumorigenic cytokine [73].

### Myc;Ptenfl tumors have shared transcriptomic signatures with human TNBC

We next used two different methods to compare the single cell transcriptomic data of our murine mammary tumors with human BC (Figure 7A). In the first method we trained a cell type classifier using public annotated human scRNA-seq breast cancer data [75] and evaluated whether the cell states observed in the Myc;Ptenfl model were consistent with those found in human disease (Figure 7A). We trained a mixture discriminant analysis classifier using the scPred R Package [76] and scRNA-seq data from 26 primary human breast tumors [75] (Supplemental figure S16). Comparison of the 28 scPred assigned cell types and the 24 original clusters found in analysis of the MycPtenfl tumors showed that most murine derived clusters were associated with a single cell type found in human data. There was high agreement for stromal cell types falling within the fibroblast, endothelial, perivascular, lymphoid and myeloid lineages with an Adjusted Rand Index (ARI) of 0.408 (Figure 7B). Notably, cluster 7 mouse fibroblast of SP showed agreement with both human CAFs: MSC iCAF-like and myCAF-like. Where mouse fibroblast cluster 17 of SR-IR showed most agreement with MSC iCAF-like and cluster 12 of SR-IP with myCAF-like (Figure 7B). Epithelial cells had weaker similarity (ARI: 0.091) between Myc;Ptenfl unsupervised clusters and scPred assigned class, likely because the original authors of the Wu data set annotated their epithelial cells with a single-cell adaptation of the PAM50 classifier rather than a data-intrinsic method such as unsupervised clustering (Figure 7B, mouse model (MM): c10, 19, 2, 8 and Human Species (HS): Cancer, LumA/B, Her2, Basal, Progenitor).

**Figure 7:**
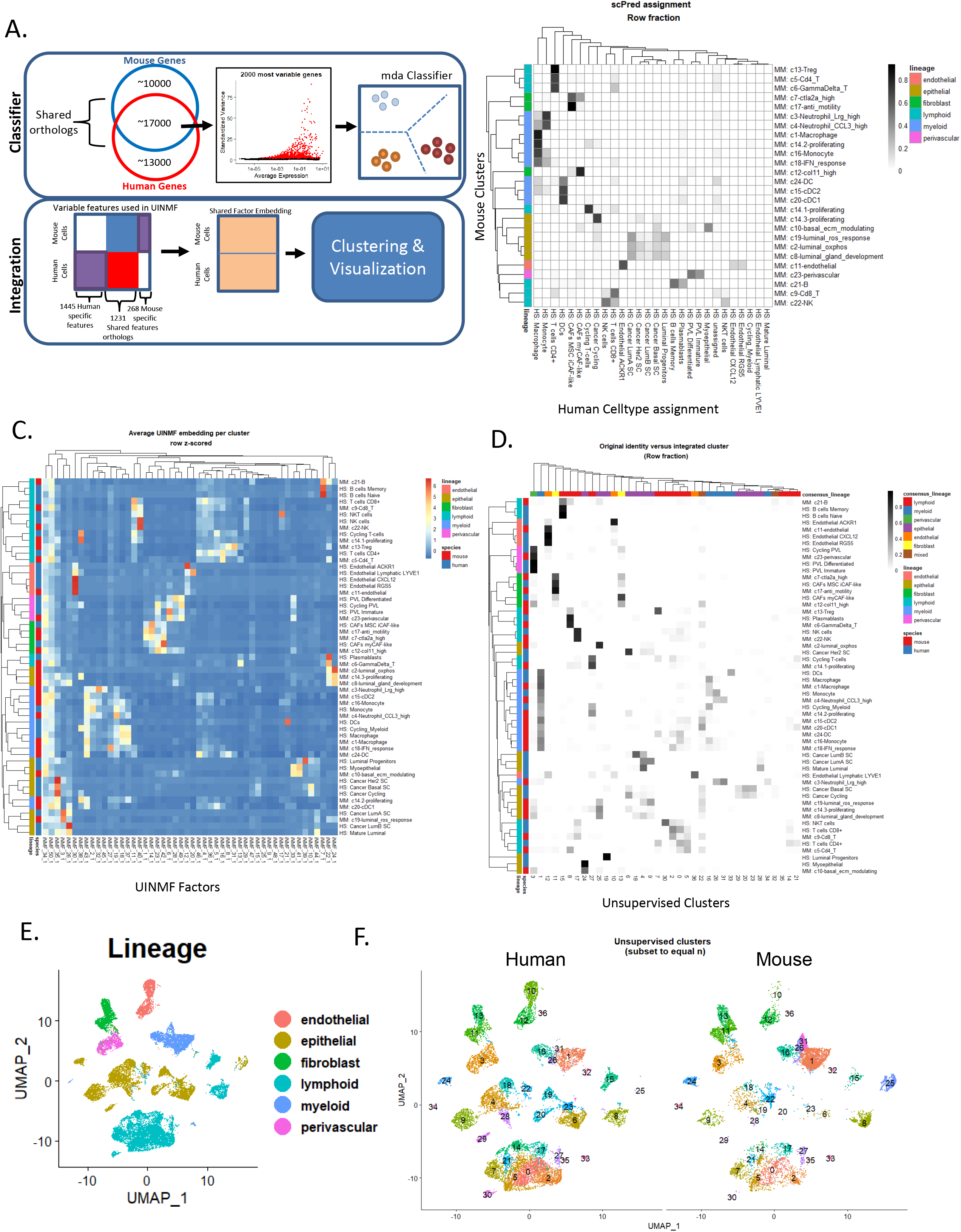
Integration of TNBC mouse model with human breast cancer scRNA-seq. **A.** Schematic showing classifier and UINMF integration. **B.** Heatmap visualizing the relationship between Myc;Ptenfl unsupervised clusters and classifier assignment from a mixture discriminant analysis based classifier built with scPred from human primary breast cancer scRNA-seq (Wu et al). Counts were row-normalized to represent the fraction of each unsupervised cluster assigned to each class. **C.** Mean UINMF embedding for each original cell type found in the Myc;Ptenfl (‘celltype_full’) and human scRNA-seq (‘celltype_minor’) data sets. **D.** Original identity (‘celltype_full’ for Myc;Ptenfl or ‘celltype_minor’ for human scRNA-seq) versus unsupervised cluster assignment. Counts were row normalized and represent the proportion of each speciesspecific cell state that was assigned to each cross-species unsupervised cluster. **E.** UMAP of UINMF integrated Myc;Ptenfl and human scRNA-seq data. UMAP was computed from 50 UINMF factors. Lineage was assigned during initial analysis of Myc;Ptenfl data (Supplemental figure 14 C) or based on author’s celltype_major classification (Supplemental dataset table 5). **F.** Unsupervised clusters of cross-species integrated data computed with the Louvain algorithm from 50 UINMF factors (Supplemental figure 16 A) Note: Point size is increased for mouse umap.

To further investigate the extent that the Myc:Ptenfl model recapitulates human disease, we integrated the Myc;Ptenfl scRNA-seq data with the primary human breast tumor scRNA-seq data (Figure 7A). Data integration and linear dimensionality reduction were performed simultaneously using Unshared Integrated Nonegative Matrix Factorization (UINMF) as implemented in the LIGER R package [77]. In total 1231 shared orthologs, 1445 human-specific genes and 268 mouse-specific genes were reduced to 50 metagene factors. The cell loadings corresponding to these 50 metagene factors were then utilized for integrated cross-species analysis. Comparison of average UINMF embedding for each cell type identified in individual species analysis indicated that the UINMF approach performed well to learn metagene factors representing shared biology conserved across species (Figure 7C). Jaccard similarity was computed to identify the overlap in the top 50 weighted genes for each factor and found that groups of factors were associated with specific cell types or lineages and shared across the two species (Supplemental Figures S16A and B).

Unsupervised clustering with the Louvain algorithm applied to the 50 UINMF factors and optimized for maximum silhouette width (supplemental figure S16C) identified 36 integrated clusters (Figure 7D). Cluster lineage was assigned for each integrated cluster by consensus of the previously assigned lineage for each cell (Figure 7D and supplemental figure S16D, with >80% agreement for lineage assignment, otherwise labelled as ‘mixed’) and only two integrated clusters represented a mixture of lineages. Notably, all 36 integrated clusters consisted of both human and mouse cells, highlighting the overlap in transcriptional signature between the Myc;Ptenfl mouse model and human BC (Figure 7D-F). Of the three cross-species clusters that were majority murine (>50% of cells in cluster came from Myc;Ptenfl model), two of them (c31, c34) consisted mostly of neutrophils (Figures 7D-F), which are known to be difficult to sequence from primary tumors due to their high RNase content [78] and were not identified in the original publication of the human data set. The remaining mouse-specific cluster 25 primarily consisted of Myc;Ptenfl cells from the epi_c2_luminal-oxphos cluster (Figures 7D-F, Supplemental Figure S16E). While this cohort of human breast scRNA-seq does not include oxphos high epithelial cells, oxphos upregulated tumors have been reported in human breast cancer and are associated with metastasis and chemoresistance [39].

## Discussion

Amplification of the *MYC* gene and loss of the *PTEN* tumor suppressor are common in human TNBC and the majority of *PTEN* loss TNBC have a copy number gain in *MYC*, which is prognostic for poor overall survival [19–21] [30]. We have generated the Myc;Ptenfl GEMM described in this work to simulate the molecular and biological complexity of MYC gain and PTEN loss observed in aggressive human TNBC. TNBCs are a heterogeneous group of tumors with one common feature: a distinctly aggressive nature with shorter overall survival and higher rates of relapse in the metastatic setting compared to other breast cancer subtypes [2–5]. A major obstacle to identifying actionable targets in TNBC is the heterogeneity of the disease, both inter and intra-tumorally. This highlights the need for robust *in vivo* models that recapitulate the spectrum of molecular and biological characteristics of TNBC, to understand the molecular basis of TNBC better and to develop effective therapeutic strategies. Our generation of the Myc;Ptenfl mouse model reveals insights into how deregulation of the *MYC* oncogene, and loss of the tumor suppresser *PTEN* can cooperate *in vivo* to generate TNBC tumors that recapitulate the heterogeneity of human TNBC subtypes as evidenced by (1) histology illustrating similar tissue architecture and cellular morphologies, (2) immunohistology displaying the presence of a similar spectrum of diverse immune cell types, (3) RNA-seq showing the precent of similar transcriptomic signatures, (4) multiplex imaging defining similar tumor and microenvironment cell phenotypes, and (5) single-cell RNA-seq revealing the presence of multiple cancer and stromal cell populations and their gene expression profiles. Myc;Ptenfl tumors recapitulate specific elements of human TNBC tumors and tumor microenvironments, such as OXPHOS high, basal-luminal mixed, and ROS high tumor cells, fibroblast heterogeneity including iCAF and myCAF populations, and immune low, immune suppressive, and immune high cell tumors corresponding to the spectrum of human TNBC. These Myc;Ptenfl tumors provide a resource for future biological studies and therapeutic testing.

An important feature of our Myc;Ptenfl TNBC model is the increased inter- and intra-tumoral heterogeneity. There are multiple Myc;Ptenfl histologic subgroups that recapitulate human TNBC subtypes and fall into two broad classes; stroma-rich, with several histological subgroup and 77% occurrence and stroma-poor with 23% occurrence. The more heterogeneous group, the stroma-rich, recapitulates the human mesenchymal TNBC subtype, marked by stromal desmoplasia, lobular 60%, squamous 15%, and metaplastic 2% IDC histology, highest expression of immune-related gene signatures and a more inflamed spatial pattern, including adjacent, peripheral and tumor-infiltrating immune cells. Gene expression analysis revealed increased hallmark of epithelial-mesenchymal-transition, tumor-promoting inflammatory response, and apoptosis. These findings are consistent with a previous human TNBC subtyping study describing mesenchymal TNBC subtypes with infiltrating immune and tumor-associated stromal cells, with elevated expression of genes involved in epithelial-mesenchymal-transition [12]. Therefore, this provides a biological rationale for using the Myc;Ptenfl-SR model in drug discovery studies requiring an infiltrating immune and tumor-associated stromal cells such as immune targeting strategies. The stroma-poor subtype recapitulates human TNBC subtypes that molecularly classifies as mixed basal like and LAR. The human data of Burstein et al. and of Ding et al. were similar to our results for mixed subtypes in that 23.7% and 33% validation samples would be classified as mixed subtypes respectively [13, 14]. This suggests that these human and Myc;Ptenfl-SP tumors have intra-tumoral heterogeneity and comprise multiple subtypes. SP is marked by solid invasive ductal carcinoma histology, high pS62MYC expression, hallmark of cellular metabolism including oxidative phosphorylation and poor prognosis. Higher MYC stability promotes an immune suppressive environment and OXPHOS signature epithelial state has been shown to enhance metastatic lethality and be associated with chemotherapy-resistant TNBC [79]. Therefore the Myc;Ptenfl-SP model can provide important insights into potential mechanisms of tumor aggressiveness and therapeutic failure that will guide future development of novel therapeutic targets in TNBC.

Interrogation of immune contexture with spatial analysis in our tumor model revealed higher total immune cells in SR, including both myeloid and lymphoid lineage cells with significant CD8 T cells in the periphery compared to SP tumors. Consistently, and similar to human TNBC, SR tumors with high tumor infiltrating lymphoid cells were associated with better prognosis whereas the SP subtype with OXPHOS metabolism, shown to be important for the production of biosynthetic intermediates necessary to support the rapid proliferation of cancer cells [39], is immune low except for macrophages and associated with high metastasis rate and poor prognosis. This type of result suggests that SP tumors would need agents to boost immune cell influx and recruitment such as chemotherapy and DC agonists, whereas SR would need agents to overcome myeloid immune suppression and re-activate T cells. SP would likely also require this approach once an inflammatory response has been evoked. The ability of the Myc;Ptenfl mouse models to represent different levels of immune activity, suggests that it may provide a robust platform for TNBC preclinical trials of newly developed immunotherapies. Currently, immunotherapy using checkpoint blockade has been shown to produce a long-lasting response in highly immunogenic cancers [80, 81]. Although breast tumors generally are not highly immunogenic, TNBC constitutes a varied spectrum of tumors with different degrees of immunogenicity that may include a more immunogenic subtype [82]. This suggests that TNBC with a higher level of lymphocyte infiltration may be more responsive to immunotherapy [81].

There is a pressing need for pre-clinical models of breast cancer which include the complete gamut of tumor microenvironment cell types to represent human disease and better assess cutting-edge therapeutic approaches. Our scRNA-seq analysis of the Myc;Ptenfl tumors revealed the extent of tumor cell and microenvironment heterogeneity and we identified discrete and transcriptionally diverse populations of epithelial (5 clusters), fibroblast (3 clusters), lymphoid (7 clusters), and myeloid cells (9 clusters). Comparison with histologically assigned tumor subtype revealed that the tumor subtype was intrinsically related to each tumor’s dominant epithelial and fibroblast cluster. We found that the SP tumors were enriched for epithelial cells with high Oxidative Phosphorylation pathway activity as well as fibroblasts with increased expression of the immune suppressive ligand Ctla2a. SR tumors with low immune presence were enriched for a heterogeneous mix of epithelial cells as well as a myofibroblasts. SR tumors with high immune presence uniquely had epithelial cells with high Reactive Oxygen Species (ROS) response pathway activity, indicating crosstalk of these epithelial cells with infiltrating immune cells [83]. ROS are also able to trigger programmed cell death (PCD) leading to apoptosis [84], a pathway that is also enriched in Stromal-rich immune-rich tumors. Furthermore, the stromal-rich immune-rich tumors were also enriched for an inflammatory CAF state with high expression of pathways related to immune activation and antimotility. Extensive comparison with a human-trained classifier and integration across species demonstrates that the cell types and states found in this murine model of TNBC are fully representative of human disease and share high transcriptional overlap with patient data [75].

In summary, our work demonstrates the important role that MYC gain and PTEN loss play in the heterogeneity and aggressive nature of TNBC. By deep analytic comparisons between our mouse model and human TNBC, we have established and characterized this preclinical Myc;Ptenfl TNBC model to resemble the morphological and transcriptional heterogeneity and key features of human TNBC disease; providing an improvement in modeling the intertumoral and TME heterogeneity seen in the TNBC patient population. These features provide insight into the molecular pathways involved in specific breast cancer subtypes and should serve as a platform for preclinical drug screening of TNBC metastasis and the testing of immunotherapies and targeted therapy strategies, as well as broad applications in basic and translational TNBC research.

## Materials and Methods

### Antibodies

HER2 (Cell Signaling #2242, 1:50); ERa (Millipore #04-227, 1:50); PR (Abcam #ab131486, 1:1000); AR (Abcam #ab47563, 1:50); cytokeratin 5 (Abcam #ab52635, 1:100); cytokeratin 14 (Covance #PRB-155P, 1:1000); pSmad3 (Abcam #ab52903, 1:100); Laminin (Abcam #ab11575, 1:50); SMA (Abcam #ab5694, 1:100); pS62 Myc rat monoclonal 4B12 [85]; Ki-67 (Abcam #15580, 1:1000); CSF-1R (Santa Cruz #sc-692, 1:500); F4/80 (Serotec A3-1, 1:200); CD11C (Cell Signaling #97585,1:100); CD4 (Cell Signaling #25229, 1:100); MHCII (eBioscience #eB14-5321, 1:100); BTK (LSBio #LS-C180161, 1:100); CD45 (BD Bioscience #550539,1:50); PDL1 (Cell Signaling #13684,1:50); CD8 (eBiosceience #14-0808082, 1:100); CD3 (Thermo #RM-9107-s, 1:300); CD207 (eBioscience #14-2073-82, 1:100); CD206 (Abcam #64693, 1:1000); B220 (BD Bioscience #550286, 1:100); RORgt (Abcam #ab207082, 1:100); Foxp3 (eBioscience #14-5773-82, 1:100); GATA3 (Abcam #ab199428, 1:100); CD11b (Abcam #ab133357,1:100); TCF1/TCF7 (Cell Signaling #2203s, 1:100); TIM3 (Cell Signaling #83882, 1:200); EOMES (Abcam #ab183991,1:1000); Granzyme B (Abcam #ab4059, 1:200); Ly6G (eBioscience #551459, 1:200); PAN Keratin (Abcam #ab27988, 1:100). CK5 (abcam, EP1601Y); S100A6 (CST, D9F9D); CD11c (CST, D1V9Y); CD103 (Biolegend, 2E7); aSMA (Santa Cruz, 1A4); EpCAM (CST, E6V8Y); CD31 (Abcam,EPR17260); ColVI (MDBiosciences, EPR17072); CD11b (Abcam, EPR1344); Ki67 (CST, D3B5); FoxP3 (Novus, NB100-39002); Vim (CST, D21H3); CD45 (CST, D3F8Q); Gal3 (Biolegend, 125408); ColIV (MDBiosciences, 203003).

### Animal studies

Rosa-LSL-Myc mice [32] were crossed with *Pten^flox/flox^* (Akira Suzuki et al. Immunity 2001) and Blg-Cre mice (gift from Owen Sansom, Beatson Institute for Cancer Research, Glasgow, United Kingdom) to generate mice that express MYC and deleted PTEN in response to Cre-mediated recombination in the mammary gland. The PTENfl and MYC;PTENfl mice investigated in this manuscript are in a FVB background and tumors were from independent animals. Tumor bearing mice were treated with Paclitaxel at dose 5mg/kg/week by intraperitoneal injection or/and DT061at dose 5mg/kg BID by oral gavage for 30 days, tumor growth was recorded on every other day by measuring the diameter in 2 cm. Tumor volume was calculated using the following formula: large diameter x (small diameter)^2^/2. If a tumor impaired an animal’s mobility, became ulcerated, or appeared infected, or a mouse showed hunched posture, the mouse was euthanized. Tumors were harvested and frozen for RNA and DNA analysis or embedded in paraffin for immunofluorescence or multiple immunohistochemistry staining.

### H&E staining, immunofluorescence and immunohistochemistry

H&E staining, immunofluorescence and immunohistochemistry were performed as described previously (Wang et al. cancer research 2010).

### Bulk RNA-sequencing and gene expression analyses

RNA-seq libraries and sequencing were performed as previously described [23]. Whole transcriptome gene expression was calculated by normalizing the read counts per transcript by the kilobases of the transcript per million mapped reads. Code used for RNA-seq analysis is available at https://github.com/zdoha/MycPtenfl-TNBC-model. For hierarchical clustering, we normalization on tumor samples using all genes, then reduced to unique gene symbols, and run gene enrichment heatmap from most variable loc and plot samples by first two principal components in R/Studio to identify tumor subgroups. Then hierarchical clustering of differentially expressed genes (enriched in luminal or mesenchymal cell lines (Prat et al., 2013) was performed using the heatmap function in R. TNBC subgrouping using the recent method developed by [14]. Using R code relies on the ggplot and reshape packages. It expects two tsv files, one for the centroids [14] and one for the RNA gene expression matrix of samples (units=TPMs). It read in those files, z-score the values across cohort, calculate the correlation to the centroids, and produce a bar plot and heatmap of the results.

### GSEA

GSEA for the murine models was performed as previously described [23].These gene sets and all GSEA results are detailed in Supplemental data set 1.

### Sequential Multiplexed IHC Staining and Analysis

Sequential IHC was performed on 5 μm FFPE sections as previously described [43, 44]. Image processing and cell quantification were performed as previously described [86, 87].

### Mice and human TMAs

The mouse Myc;Ptenfl TNBC TMA was generated by marking tumor regions of interest on FFPE blocks and punching 1.5 mm cores using TMA Master II (3DHistech, Hungary) for drilling recipient block and MTA-1 (Estigen Tissue Science, Estonia) for tissue coring. The tumors in the human TMAs were determined to be TNBC by pathologist (MES) evaluation of IHC staining for ER, PR, and HER2. TMA101 and TMA11-4-09 were constructed from surgically resected primary tumor samples from patients with breast cancer diagnosed at Vanderbilt University Medical Center. One millimeter tumor cores (two per surgical specimen) were punched from representative areas containing invasive carcinoma selected by a pathologist. Clinical and pathological data were retrieved from medical records under institutionally approved protocols, IRB# 030747 and 130916, for patients in TMA101 and TMA11-4-09, respectively.

### Mice and human TNBC TMA H&E histologic/morphologic analysis

Pipeline for generating morphological feature representation using a variational autoencoder (VAE) to compare tissue microarray (TMA), from our TNBC mouse model with TMA from Human TNBC. First, raw Hematoxylin & Eosin (H&E) stained TMA images (92 TMA cores from mice TMAs and 172 cores from human TMAs) were pre-processed to account for between-sample intensity variation. H&E pixel intensities were normalized using the Reinhard method [88] where background pixels are excluded from intensity distributions. One subset of human cores was used as the target distribution to which each other dataset was normalized [89]. For parameter selection and optimization, we explored H&E tile size of 256×256, 512×512, 1024×1024, 2048×2048 (pixels), and latent feature vector dimension of 32, 64, 128, 256, 512 to identify meaningful representations of the image dataset. We found that 1024×1024 tile size and a feature vector dimension of 64 captured meaningful histological features based on visual evaluation and yielded the lowest reconstruction losses. Tiles from both mice and human TMAs are used to train a VAE, then a latent encoding vector is computed for each tile. Tiles are compared using UMAP embedding and k-means clustering analysis of the latent features. Density functions for all human and mice tiles are calculated within the 2-dimensional UMAP space to visually compare overlap in embedding space. K-means clusters (n=8) are computed using latent features and projected into UMAP space for visualization. The relative abundance of human and mice TMAs are calculated for each cluster using the ratio of tiles in a cluster to total tiles from the given TMA source.

### Cyclic Multiplexed-Immunofluorescence (cmIF) and single-cell multiplexed analysis

Cyclic immunofluorescence, image processing and Cyclic IF analysis were performed as previously described [63].

Cyclic IF Analysis: Code used for analysis is available at https://github.com/engjen/MYC-PTENfl-mouse. Single-cell mean intensity values of autofluorescence-subtracted images were selected from either nucleus or cytoplasm masks for each marker, depending on expected intracellular distribution. Areas of floating tissue that created bright imaging artifacts and air bubbles that created dark artifacts were manually circled using the napari image viewer and excluded. Background subtraction was performed on markers with high background: CD31, CD45, CD8, ColIV, FoxP3, CD103, CD11b and CD11c. Filtered, background subtracted data were clustered with scanpy [90]. Two morphology features (nuclear area and nuclear eccentricity) and 20 markers were used for clustering. A Umap embedding was generated using 15 neighbors, and the Leiden algorithm was used for unsupervised clustering (resolution 0.6), resulting in 20 cell types. Inspection of cluster results in images revealed that 3 of the clusters were due to imaging artifacts and excluded. The remaining clusters were evaluated on the images and annotated. Endothelial cells were separated from mixed endothelial/immune and endothelial/fibroblast clusters by manually gating based on CD31 expression.

We then performed manual gating to verify our annotated-cluster cell type calling. A threshold was set for each gating marker based on positive pixel patterns in images. Endothelial cells were defined as CD31+. Epithelial cells were positive for 1 or more of Ecad, EpCAM and CK5 and CD31-. Immune cells were CD45+ CD31- and epithelial marker negative. Stromal cells were all non-endothelial, non-epithelial, non-immune segmented nuclei. Three cores had pMYC positive tumor cells negative for Ecad, EpCAM and CK5 (I11, G09 and H11). In these tissues, tumor cells were defined as any cells negative for all stromal and immune gating markers (CD31, CD45, Vim, aSMA, ColIV, ColIV, Gal3).

Cell frequencies were calculated for gated epithelial, immune and stromal cells in each tissue core. Endothelial cells were rare and added to “stromal” cells for subtyping. Tissues were clustered with the Leiden algorithm, resolution = 0.5, resulting in 6 subtypes. Subtypes were annotated as stroma-rich, immune-rich (clusters 2 and 4), stroma-rich, immune-poor (cluster 0) and stroma poor (cluster 1, 3 and 5). Chi-squared analysis was performed on cell frequency subtype versus histology subtype (i.e. stroma-rich and stroma-poor based on H&E evaluation). Most of the cyclic IF stroma poor samples were also stroma poor by histology (6/9). Discrepancies between CyCIF subtypes and histology subtypes were examined (Supplemental Figure S17) and found to be a result of tumor-adjacent stroma, lymphocytes in the tumor core or significant nuclei-free areas of tissue.

To compare expression of markers in each subtype, mean marker expression was calculated in each tissue and compartment (epithelial or stromal, i.e. immune, endothelial and non-immune stromal cells). Distributions were visualized as boxplots and the Kruskal Wallis H test of Mann Whitney U test, implemented in scipy [91], were used to test for significant differences in the median expression of markers between groups. To compare detailed cell types (14 annotated cell types from Leiden clustering, described above) between subtypes, the frequency of each cell type in each subtype was calculated and displayed as a bar plot. Samples were also clustered hierarchically based on the z-score of cell abundances.

For human tissue samples, we obtained a publicly available multiplex ion beam imaging (MIBI) dataset [64] at https://github.com/aalokpatwa/rasp-mibi. We used scanpy for single-cell clustering. Thirty-two markers were used for generating a Umap embedding with 15 neighbors. Unsupervised clustering was then performed with the Leiden algorithm (resolution = 0.6) resulting in 24 cell types. These were annotated as epithelial, immune or stromal. Inspection of cluster results versus images revealed that some clusters contained mixed immune and stromal cells. Therefore, epithelial clusters were used to define epithelial cells, and CD45 and CD31 were used to manually gate immune and endothelial cells within the stromal/immune clusters. Tissues were clustered on cell type frequencies with the Leiden algorithm, resolution = 0.1, resulting in 3 subtypes, annotated as stroma-rich, immune-rich (cluster 0), stroma-poor (cluster 1) and stroma-rich, immune-poor (cluster 2). Mean marker expression was calculated in each tissue and compartment, visualized and statistically evaluated as described for mouse data, above. For survival analysis, the lifelines [92] python software was used. Kaplan-Meier estimates were generated and plotted for overall survival and recurrence-free survival. The log rank test was used to test for significant survival differences between the subtypes.

### scRNA library preparation and sequencing

Single-cell suspensions of Myc;Ptenfl breast cancer tissues were obtained by enzymatic digestion. Tissue was manually minced using scissors, followed by a 30-60 min enzymatic digestion with 2.0 mg/ml collagenase A (Roche), 1.0 mg/ml Hyaluronidase (Worthington), and 50 U/ml DNase I (Roche) in serum-free Dulbecco’s modified eagles medium (DMEM) (Invitrogen) and Rock inhibitor at 37C using continuous stirring conditions. Single cell suspensions from tumor digests were prepared by passing tissue through 40-mm nylon strainers (BD Biosciences). Single cell suspensions from individual tumors were then labeled with hashtag oligonucleotides following the manufacturer’s protocol (TotalSeq B0301-B0306, Biolegend). Each individual tumor sample was counted and then pooled at an equal cell ratio before being split into two replicates for library preparation with the Chromium Single Cell 3’ V3 (10X Genomics) following the manufacturer’s protocol with a targeted recovery of 20,000 cells per library. Libraries were sequenced on an Illumina NovaSeq. BCL files were converted fastq format with bcl2fastq2 (Illumina), and then aligned to mouse genome build mm10-2020-A (10X Genomics) using Cellranger (10X Genomics, version 6.0.2).

### scRNA-seq data processing

The R package SoupX was used to load the UMI gene count matrix for each library and correct for ambient mRNA contamination using default parameters(Young and Behjati, 2020). The corrected UMI gene count matrix was then converted to Seurat Object format using Seurat (v4.1.0) and the paired HTO count matrix was added. HTO demultiplexing was performed using the *HTODemux* function of Seurat with parameters: (kfunc = ‘clara’, positive.quantile = 0.95) (Satija *et al*., 2015; Butler *et al*., 2018; Stuart *et al*., 2019; Hao *et al*., 2021). Doublets were identified within each library using the R package DoubletFinder (v2.0.3) with a presumed Poisson doublet rate of 0.075 and 10 principal components(McGinnis, Murrow and Gartner, 2019). Only cells with greater than 250 unique genes expressed, less than 25% mitochondrial RNA and assigned as a ‘Singlet’ via DoubletFinder were retained for analysis.

### Myc;Ptenfl scRNA-seq normalization, integration, clustering, and differential expression analysis

UMI counts were log normalized and scaled without centering using the R package Seurat (v4.1.0). The top 2000 variable features were identified using the VST method using the R package Seurat. iNMF integration was performed directly on the Seurat object accessing the rliger package (v1.0.0) via the SeuratWrapper package (v0.3.0). OptimizeALS was run with the parameters: (k = 50, lambda = 5, nrep = 5, split.by = ‘library_id’). RunQuantileNorm was performed with the same split.by setting. UMAP visualization was performed using the resultant 50 iNMF factors. Unsupervised clustering was performed using the Leiden algorithm with a resolution of 0.45, which corresponded to the point of diminished returns for increased silhouette width as estimated by the ‘approxSilhouette’ function of the R package Bluster (v1.2.1).

### scRNA-seq differential expression and enrichment of gene ontology

Differential expression analysis was performed using the Wilcoxon rank-sum test via Seurat’s ‘FindMarkers’ function, and significantly differentially genes for any cluster had at least a log2 fold-change of 0.5 and Bonferroni corrected p value below 0.05. The R package ClusterProfiler (v4.0.5) was used to identify enriched gene ontologies from significantly upregulated genes with parameters: (ont = ‘ALL’, pAdjustMethod = ‘BH’, pvalueCutoff = 0.01, qvalueCutoff = 0.05).

### Cross-species classifier

Orthologous genes found in both the Wu et al dataset and Myc;Ptenfl scRNA-seq data were found using the ‘convert_mouse_to_human_symbols’ function in NicheNetR (v1.0.0). The Wu dataset was subset to only include orthologous features, and a classifier was trained using the mixture discriminant analysis (MDA) model of scPred (v1.9.2). The classifier was then applied to the Myc;Ptenfl data with a threshold of 0.55. Adjusted Rand Index was computed with the R package aricode (v1.0.0) and used to compare the original labels derived from unsupervised clustering and transferred classes from scPred classifier.

### Cross-species scRNA-seq data integration and analysis

Orthologous genes found in both the Wu et al dataset and Myc;Ptenfl scRNA-seq data were found using the ‘convert_mouse_to_human_symbols’ function in NicheNetR (v1.0.0) and the mouse features were updated to human symbols. The UMI count matrices for mouse and human datasets were loaded with rliger (v1.0.0) and log normalized. Variable genes were found with the ‘selectGenes’ function of rliger using parameters:(var.thres = 0.3, unshared = TRUE, unshared.thresh = 0.3, unshared.datasets = list(1,2)), and 1231 shared features, 268 mouse features and 1445 human features were selected. Variable features were scaled without centering in rliger, and then the optimizeALS function was run to perform matrix factorization using parameters: (lambda = 5, use.unshared = TRUE, thresh = 1e-10, k = 50, nrep = 5). The rliger ‘quantile_norm’ function was then used to build the shared factor graph and normalize with parameter: (ref_dataset = ‘human’). The liger object was then converted to Seurat format with the rliger function ‘ligerToSeurat’. The cell loadings from all 50 UINMF factors were used for UMAP embedding and unsupervised clustering. Unsupervised clustering was performed using the Louvain algorithm as implemented in Seurat, and a resolution of 0.6 was selected as it optimized for maximum mean silhouette width as estimated with the ‘approxSilhouette’ function of the R package Bluster (v1.2.1). Unsupervised clusters were assigned lineage if there was at least 80% agreement of prior lineage annotation of the cluster’s constitutive cells, otherwise they were labeled ‘mixed’.

### Study approval

All protocols for mouse experiments described in this study were approved by the Oregon Health & Science University Animal Care and Use Committee protocol # IP00001014, Portland, OR.

### Statistics

Spearman correlation coefficient was used to assess correlations of percentages and densities among tumor samples lineages. An unsupervised hierarchical clustering was performed with Ward’s minimum variance method (“hclust” from “R”). All statistical calculations were per-formed by R software, version 3.5.2 (https://www.r-project.org). P < 0.05 was considered statistically significant.

Statistical analysis was performed using GraphPad Prism software. Measurements are presented with sample n and mean +-SD or SEM as indicated in figure legends. An unpaired two-tailed Student’s T test was used throughout to compare two groups. A base P value of <0.05 was considered statistically significant.

## Acknowledgment

We thank Zuzana Tatarova’s and Dylan Blumberg’s work for antibody labeling and validation for the CycIF staining. RNA-sequencing was performed in the Massively Parallel Sequencing Shared Resource at OHSU. Some of the embedding and sectioning of tissues was performed by the OHSU Histopathology Shared Resource, and some of the imaging was performed by the Advanced Light Microscopy Core. The OHSU Shared Resources are supported by the Knight Cancer Institute through NIH P30 CA69533. This project was funded in part by the OHSU Knight Cancer Institute. Funding for these studies include U54 CA209988 to JWG and RCS, R01 CA186241, R01 CA129040, U01 CA224012, Prospect Creek Foundation, and Brenden-Colson Center foundation to RCS; NIH P50 CA098131 and Komen SAB210301 to JAP.

## Author Contributions

ZOD, XW, and RCS designed the study. MM examined the METABRIC dataset of primary breast cancer. XW performed all animal studies and ZOD assisted with monitoring mouse tumors for single-cell suspensions. ZOD and CP perform the bulk RNA-Seq analysis and clustering and ZOD generated the data presentation and performed all GSEA analyses and human correlation. LT, ENK, YHC performed the tissue microarray (TMA) histological and morphological analysis, and write up. XW, CB, NK and LMC performed or supported the mIHC staining and analysis; ZOD assisted with data presentation. JAP, MES, TW and XW generated the human and mouse tissue microarrays, JWG and KC directed CycIF workflow, SK, EB and JE performed CycIF staining, image processing and multiplex imaging analysis for the TMA. ZOD, NC, CJD, XL, LMH and GM performed or supported the scRNA library preparation and analysis; NC performed the scRNA-Seq analysis, clustering and cross-species scRNA-seq data integration and write up. ZOD and RCS wrote the manuscript. All authors reviewed and edited the manuscript.

## Supplemental Datasets

Supplemental Dataset 1: GSEA analysis of RNA-seq

Supplemental Dataset 2: Human and Myc;Ptenfl mice TMA histologic features

Supplemental Dataset 3: mIHC Antibody table

Supplemental Dataset 4: mTMA-Marker Table

Supplemental Dataset 5: Cell type_markers used in scRNA-seq analysis

## Data availability

The data generated in this study will be available to public in Gene Expression Omnibus (GEO) on October 17, 2022

