## supplemental figures for "A Novel Mouse Model that Recapitulates the Heterogeneity of Human Triple Negative Breast Cancer"

**Supplemental Figure Legends:**

**Supplemental Figure S1. Supplemental Figure S1. A.** Examining the METABRIC dataset (Curtis et al., 2012), with over 2,400 primary breast cancer tumors for low-level gain and high-level amplification of c-Myc occurs. **B**. Overall survival of MYC CNAs**. C.** Immunohistochemical staining for the indicated proteins in SR tumors (N=20/20 positive SMA, pSMAD3, Cytokeratin 5 (KRT5) , high collagen) and SP tumors (N=10/10 negative SMA, pSMAD3, low KRT5, low collagen). Scale bars = 100 (SMA), 50 (pSMAD3), 100 (K5), 200 (Trichroma) µm. **D**. On the left; Immunohistochemistry(IHC) staining with anti-phospho-MYC in stromal rich vs stromal poor tumors, Scale bar= 50µm. On the right; the percentage of pho-MYC positive cells were analyzed in stroma rich tumors (N=4) compare to stroma poor tumors (N=4). Scale bars = 50 µm **E.** total metasteses rate for Ptenfl, Myc;Ptenfl-SR and Myc;Ptenfl-Sp; Pten;Blg 3/19=16%. Myc;Ptenfl SR: Macro 9/23=39%; Micro 3/23=13%. Myc;Ptenfl SP: Macro 5/15=33.3%; Micro 4/15= 26.7%.

**Supplemental Figure S2: A**. Plot by first 2 principal components; SR (red), SP (blue). **B.** Normalize expression values used to create heatmap from most variable loci from 1200 most variably expressed genes, recapitulates 2 subgroups. **C.** Gene set enrichment analysis (GSEA) comparing SR tumors to SP tumors from MYC-PTENfl mice. **D.** A heatmap comparing expression of 43 lineage-correlated genes in tumors from the 2 groups of Myc;Ptenfl, including 14 luminal genes, 15 Basal genes and 14 mesenchymal genes. **E.** On the top; Immunohistochemistry (IHC) staining with anti-AR in stromal rich vs stromal poor tumors, Scale bar= 100µm. On the bottom; the percentage of AR positive cells were quantified in stroma rich tumors (N=5) compare to stroma poor tumors (N=5).

**Supplemental Figure S3: Myc;Ptenfl tumors show heterogeneous triple-negative expression at an early stage. A.** Multiple histologic subtypes present early in small tumors (dimeter <3mm) in 32 MYC-PTENfl mice. Top panel Scale bar =1mm, bottom panel scale bar =200um **B.** Pie chart of the histological frequency subtypes in 32 small tumors from MYC;PTENfl mice. **C.** On the left, Immnuhistochemistry staining of MYC;PTENfi mice early stage tumors with anti-Era, anti-HER2, and anti-PR in papillary (N=10/10 negative), squamous (N=10/10 negative), lobular (N=10/10 negative) subtype tumors and solid tumors (N=10/10 negative); postivie Era and PR staining in adjacent normal ducts indicated below. On right, Immnuhistochemistry staining of MYC;PTENfl mice early stage tumor with the basal marker KRT14 in papillary (N=10/10 positive), squamous (N=10/10 positive), lobular (N=10/10 positive) subtype tumors and solid tumors (N=10/10 negative). Scale bars = 100 µm.

.

**Supplemental Figure S4: A.** Determine tumor border by CD45 marker**. B.** Determine tumor border by PANCK expression. **C,D,E.** The distribution of each lineage of immune population in stromal rich compare to stromal poor tumor; core, border periphery, respectively **F.** Total Treg $CD4^+$ cells with $Foxp3^+$ population in stromal rich tumor compare to stromal poor tumor. **G.** Non-Treg $CD4^+$ cells in the stromal poor tumors compare to stromal rich.

**Supplemental Figure S5: Histopathological features space projection of tile images from human and mice TMAs. A.**The learned VAE feature representations for each of the tiles in the TMA dataset are projected into two dimensions with the UMAP embedding. **B.** The learned VAE feature based on human subtype information (TNBC and non TNBC core in the human TMA).

**Supplemental S6: Mice TNBC showed many well-known human TNBC histo-morphological features. A-D.** Mice TNBC showed various stages of breast cancer from IDC with low-grade nuclear feature (A) to high-grade nuclear feature (B), sarcomatous transformation (C), and solid growth pattern (D). **E-H**. In addition, mice TNBC characteristically showed squamous-myoepithelial differentiation. The transition between squamous cells (yellow arrow) and myoepithelial cells (white arrow, E, medium magnification, F, high magnification). Luminal epithelial (red arrow) and myoepithelial (white arrow) proliferation (G) . Squamous cell carcinoma (yellow asterisk) along with sarcomatous cancer (white asterisk, H ). **I.** Stromal cell with inflammatory cell infiltrations, **J.** Geographic necrosis (white asterisk), **K**. Clear cell change, **L**. Neuroendocrine differentiation with thick trabeculae. Scale bars = 22 μm, 22 μm, 22 μm, 44μm, 44μm, 22μm, 22μm, 44μm, 44μm, 44μm, 22μm, 44μm respectively. **M**. Frequency of human TNBC characteristic histologic features in Myc;Ptenfl mice TNBC model.

**Supplemental Figure S7: Gating and unsupervised clustering-based cell type definition in mouse and human tissues. A.** UMAP projection of single cells in mouse TMA, colored by expression of cell type specific markers. **B.** Manual gating of mouse epithelial, immune and stromal cell types projected on the UMAP. **C.** Unsupervised clustering with the Leiden algorithm resulted in 21 cell types in mouse tissues, 17 of which were subsequently annotated and 4 of which were imaging artifacts. **D.** UMAP projection of single cells in human TNBC, colored by expression of cell type specific markers. **E.** Manual gating of human epithelial, immune and stromal cell types projected on the UMAP. **F.** Unsupervised clustering with the Leiden algorithm resulted in 25 cell types in human tissues.

**Supplemental Figure S8. Cell frequency-based subtyping in mouse and human tissues. A**. UMAP embedding of mouse tissues based frequencies of epithelial, immune and stromal non-immune cells in each TMA core (k-nearest neighbors k=5). **B.** Histology-based subtypes projected on same UMAP in (A). **C.** Unsupervised clustering of mouse tissues with the Leiden algorithm (resolution 0.5) resulted in six clusters, annotated as three subtypes: stroma-poor (SP), stroma-rich, immune-rich (SR_IR) and stroma-rich immune-poor subtype (SR_IP). **D.** UMAP embedding of human tissues based frequencies of epithelial, immune and stromal non-immune cells in each ROI (k-nearest neighbors k=5). **E.** Unsupervised clustering of TNBC patients using the Leiden algorithm (resolution 0.1) resulted in three clusters annotates as: stroma-poor (cluster 1), stroma-rich, immune-rich (cluster 0) and stroma-rich immune-poor subtype (cluster 2). **F.** Kaplan-Meier curves of recurrence-free survival in cell frequency-based subtypes in human TNBC, log-rank p-value = 0.63. **G.** Kaplan-Meier curves of overall survival in cell frequency-based subtypes in human TNBC. Log-rank p=0.046.

**Supplemental Figure S9. Histological and image-based validation of cell frequency subtypes. A.** Hierarchical clustering of mouse samples based on cell type frequency in each tumor. Row annotations show Leiden clusters, annotated subtypes, and histology subtypes. **B.** Hierarchical clustering of mean cell type frequency of six Leiden clusters from (A). Row annotations show meta-clusters/annotated subtypes: Very stroma poor (SP+, red), stroma poor (SP, orange), stroma-rich, immune-rich (SR_IR, blue) and stroma-rich, immune poor (SR_IP, green). **C.** Three annotated subtypes based on cell type frequency (y-axis) versus histology subtypes (x-axis), Chi-squared p=0.26. **D**. Four annotated subtypes based on cell type frequency (y-axis) versus histology subtypes (x-axis), Chi-squared $p < 0.0001$. **E.** Representative three-color MIBI images of a 800 X 800 µm region-of-interest from stroma poor (left) and stroma rich, immune rich subtypes (right) of human TNBC. **F.** Gated cell type spatial distribution of tissues in (E). Orange: epithelial, pink: immune and green/blue: stromal cells.

**Supplemental Figure S10: A.** Stromal expression of myeloid-lineage (CD11b), activated fibroblast/pericyte (alpha-SMA), mesenchymal (Vim) markers, and epithelial nuclear eccentricity and epithelial pMYC expression in mouse subtypes. **B.** Stromal expression of lymphocyte (CD20, CD3, CD8, CD4, CD56, CD138), monocyte (CD11c, CD63), immune checkpoint (PD-L1, Lag3, IDO, PD1), memory (CD45RO), antigen presentation (HLA-DR), activated fibroblast/pericyte (SMA), mesenchymal (Vimentin), and macrophage (CD68) markers in human subtypes. **A-B.** P-values determined by Kruskal–Wallis H test. **C.** Epithelial expression of phospho-MYC, basal/myoepithelial (CK5, alpha-SMA), mesenchymal (S100A6) and epithelial nuclear eccentricity in mouse histological subtypes. P-values determined by Mann–Whitney U test.

**Supplemental Figure S11: Marker expression in human cell types defined by unsupervised clustering and hierarchical clustering of mouse and human tissues. A.** Six-color overlay of intratumoral and stromal heterogeneity in mouse tissue core. **B.** Unsupervised-clustering based detailed cell types reflect the spatial heterogeneity observed in (A).**C.** Mean marker intensity in each annotated cell type defined by unsupervised Leiden clustering in human TNBC tissues. Row annotations show cell lineage: epithelial (light orange), immune (blue) or non-immune stromal (green) endothelial (purple). **D.** Hierarchical clustering of human samples based on detailed cell types. Heat map column colors: stroma-poor (SP, orange), stroma-rich, immune-rich (SR_IR, blue) and stroma-rich-immune-poor (SR_IP, green). **E.** Bar plot of frequency of each cell type in human subtypes.

**Supplemental Figure S12:** H&E staining for the 11 Myc;Ptenfl tumor samples used for scRNAseq analysis. A. stromal rich histology for 7 out of 11 Myc;Ptenfl tumors. B. stromal poor solid histology for 4 out of 11 Myc;Ptenfl tumors. Scale bar = 100 µm.

**Supplemental Figure S13: scRNA-seq quality control. A.** Violin plots showing QC statistics for each individual Myc;Ptenfl tumor. Boxplots line represents median value, boxes are inner quartile range (IQR) and whiskers extend to the furthest point less than 1.5*IQR from median. **B.** Number of cells passing each filtering threshold per tumor. **C.** Table summarizing the number of cells, mean number of UMI recovered, mean number of unique genes, and mean percent of UMI contributed from mtRNA for each individual tumor.

**Supplemental Figure S14: iNMF data integration and clustering. A.** UMAPs computed from: 50 principal components with no integration, 50 harmony integrated principal components, 50 iNMF factors computed with Rliger. Integrations were performed across scRNA-seq libraries to remove technical noise while retaining biological heterogeneity. **B.** Clustering quality results computed on the 50 iNMF factors across a range of Leiden resolutions (0.1 – 1.5). Mean RMSD and mean Silhouette Width were computed for each resolution using the R package Bluster. A resolution of 0.45 was selected as optimal and used for downstream analysis as it represented a plateau in silhouette width. **C.** Dotplot showing the average expression of select lineage related markers for each unsupervised cluster. **D.** UMAP of Cluster 14 (proliferative cluster) computed on 5 harmony components. Harmony was used to integrate the subset of cluster 14 across sequencing libraries, and 5 harmony components were chosen based on elbow plot for UMAP visualization and unsupervised clustering. **E.** Dotplot of the same lineage markers as C for cluster 14 subclusters shows that cluster 14 includes proliferating cells from three lineages (lymphoid, myeloid, and epithelial).

**Supplemental Figure S15: iNMF data integration and clustering. A.** Dotplot showing up to the top 3 differentially expressed genes for each unsupervised cluster when compared to all other clusters. **B.** Heatmap of the mean iNMF cell embedding for each lineage. Hierarchical clustering was performed on both rows and columns and is represented by dendrograms. **C.** Heatmap of the mean iNMF cell embedding for each unsupervised cluster. Hierarchical clustering was performed on both rows and columns and is represented by dendrograms. **D.** On the left: Barplot showing mean celltype frequency within lineage for each MycPtenfl tumor subtype. Error bar indicates SEM. On the right: Relative proportion of each unsupervised cluster for each tumor, color coded by Myc;Ptenfl subtype. **E.** Mean cluster expression of Epithelial biomarker genes associated with basal, luminal, or mesenchymal cell states. Expression scaled to represent z scores for epithelial cells.

**Supplemental Figure S16: Cross-species scRNA-seq comparisons. A.** Jaccard similarity index computed for the top 50 weighted shared features for each UINMF factor. **B.** Mean UINMF cell embedding for each lineage computed for each factor. UINMF rows are ordered the same as figure A. **C.** Clustering quality results computed on the 50 UINMF cell embeddings across a range of Louvain resolutions (0.3 – 1.2). Mean RMSD and mean Silhouette Width were computed for each resolution using the R package Bluster. A resolution of 0.6 was selected as optimal and used for downstream analysis as it was the maximum mean silhouette width. **D.** Fraction of each lineage (assigned during single-species analysis) for each cross-species UINMF integrated cluster. Clusters with >80% consensus were assigned their respective majority lineage, and clusters without consensus were labeled as ‘mixed’. **E.** Distribution unsupervised clusters present in each sample. Rows represent a unique sample (either patient from Wu et al data, or individual tumor from Myc;Ptenfl model) and columns are unsupervised clusters computed on the UINMF integrated data. Rows are annotated by species and tumor subtype (ER+, HER2+, TNBC for human data or SR-IR, SR-IP, or SP for Myc;Ptenfl data). Columns are annotated by consensus lineage as assigned in figure D.

**Supplemental Figure S17: Examples of discrepancies between CyCIF and histology subtypes.** Central heatmap of frequency of epithelial, immune and non-immune stromal cells in

CyCIF data with rows annotated by histology subtypes. Black lines indicate examples of agreement of disagreement between CyCIF and histology, including: 1. The edge of the core includes some tumor adjacent stroma, so the SP histology gets called as SR by CyCIF (cores D7, B7). 2. The core has acellular areas with no or few cells segmented, so SR histology gets called as SP by CyCIF (Core B9). 3. There are a number of intratumoral lymphocytes that are hard to distinguish in H&E (without CD45), so SP histology becomes SR, immune-rich with CyCIF characterization (cores A5, H11). 4. Most stroma poor by histology are also stroma poor by CyCIF (core G8). Scale bar in core B9 is 130 μm.

#### Supplemental Figure S1

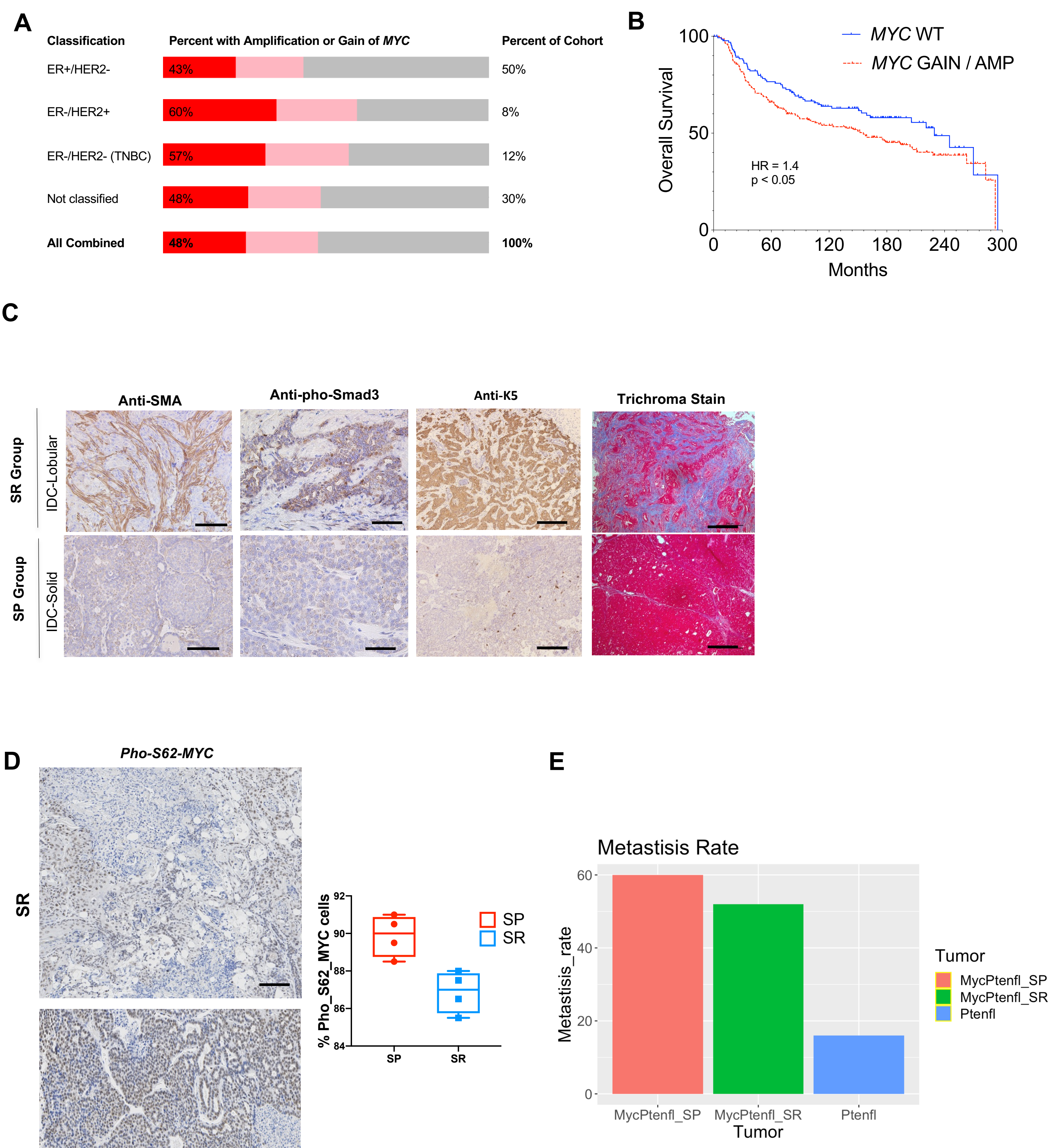

**Supplemental Figure S2**

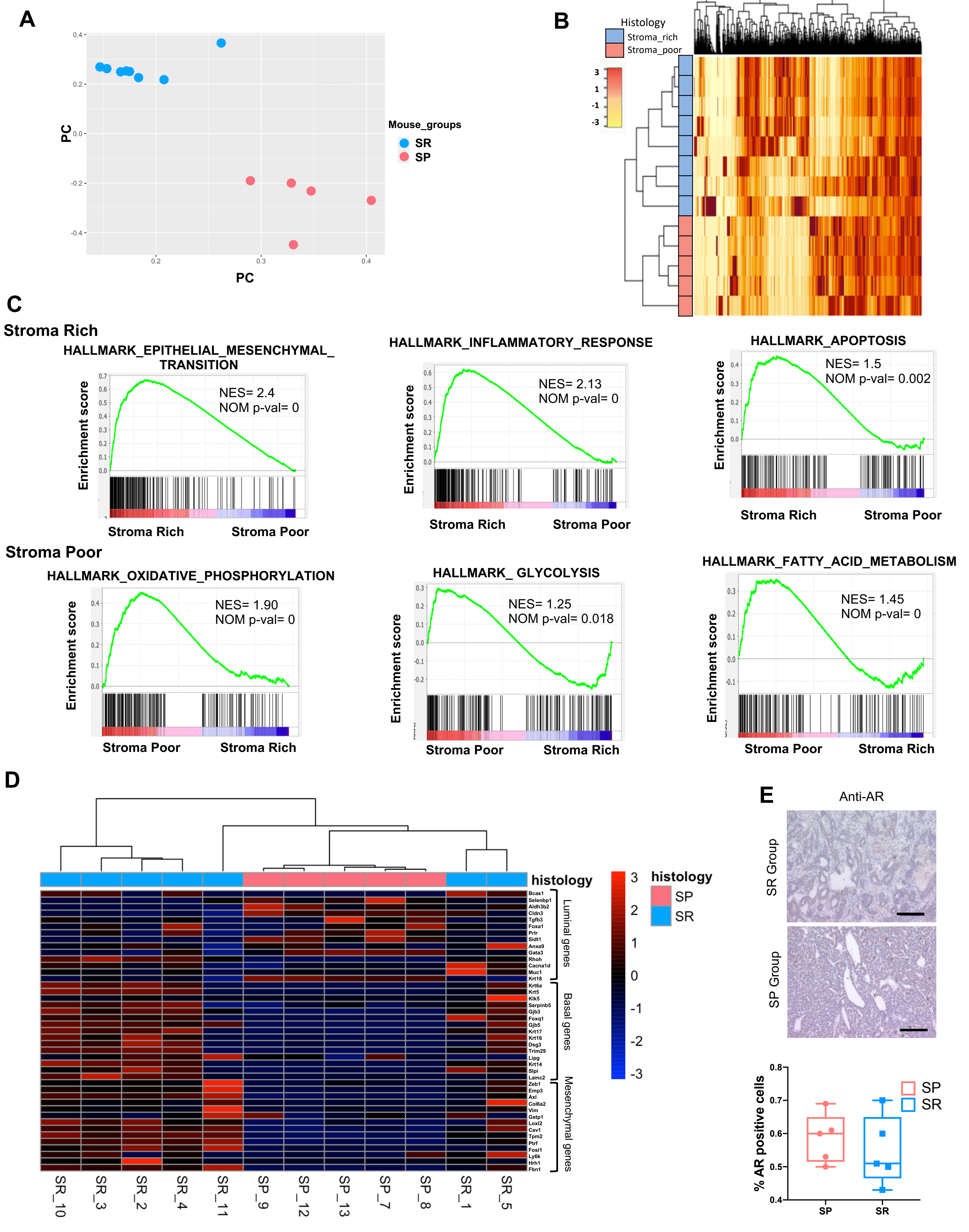

### Supplemental Figure S3

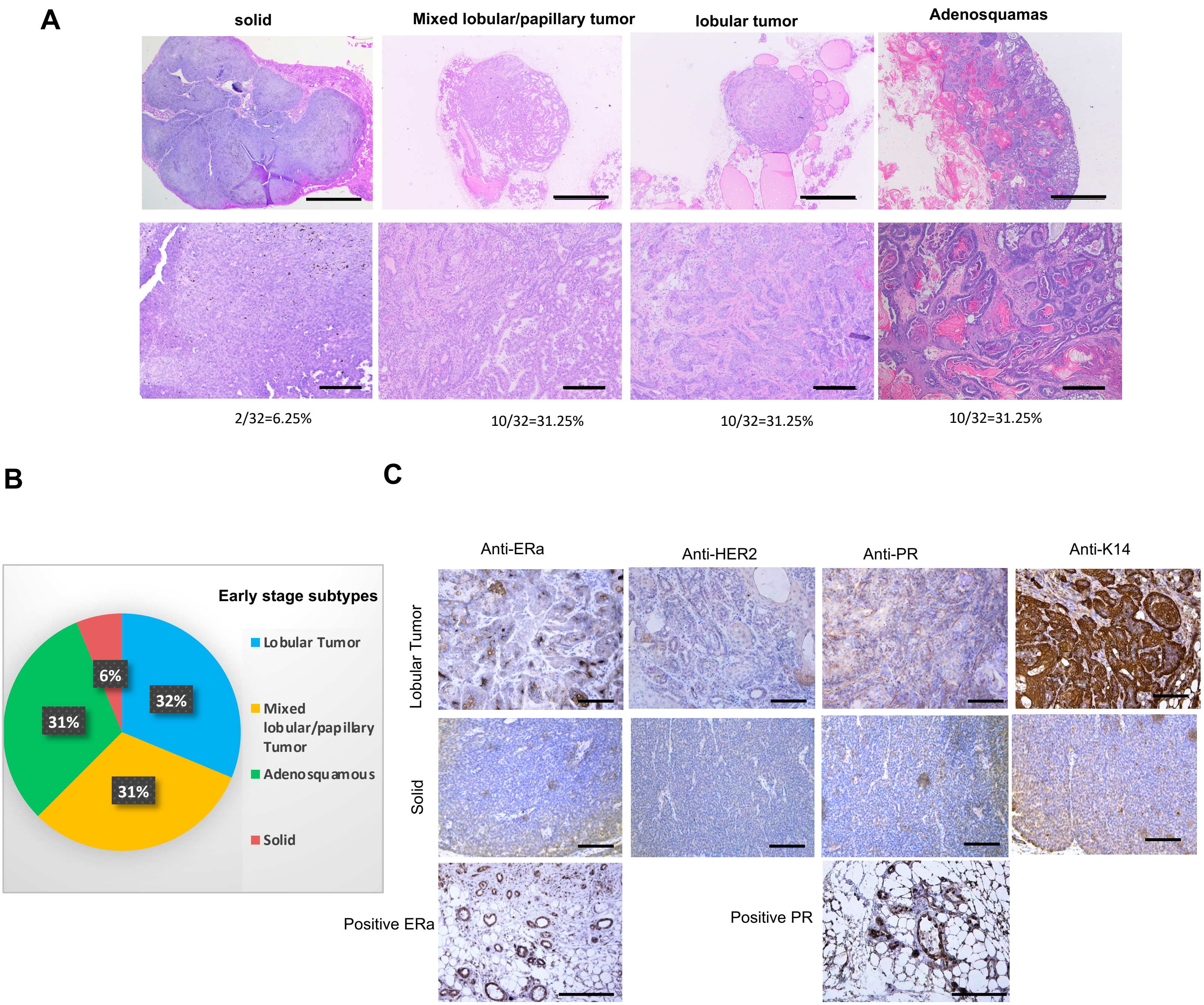

### Supplemental Figure S4

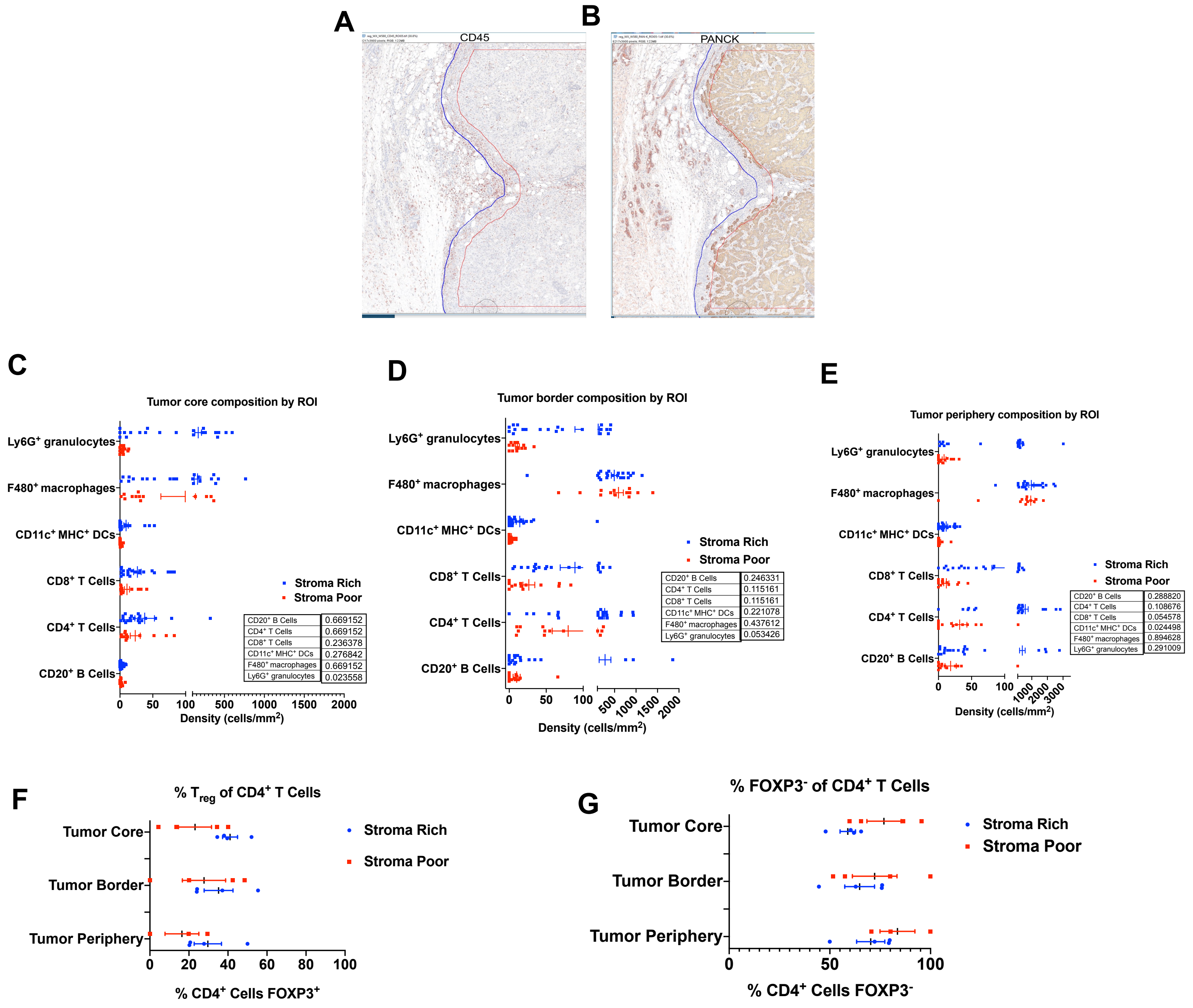

**Supplemental Figure S5**

A

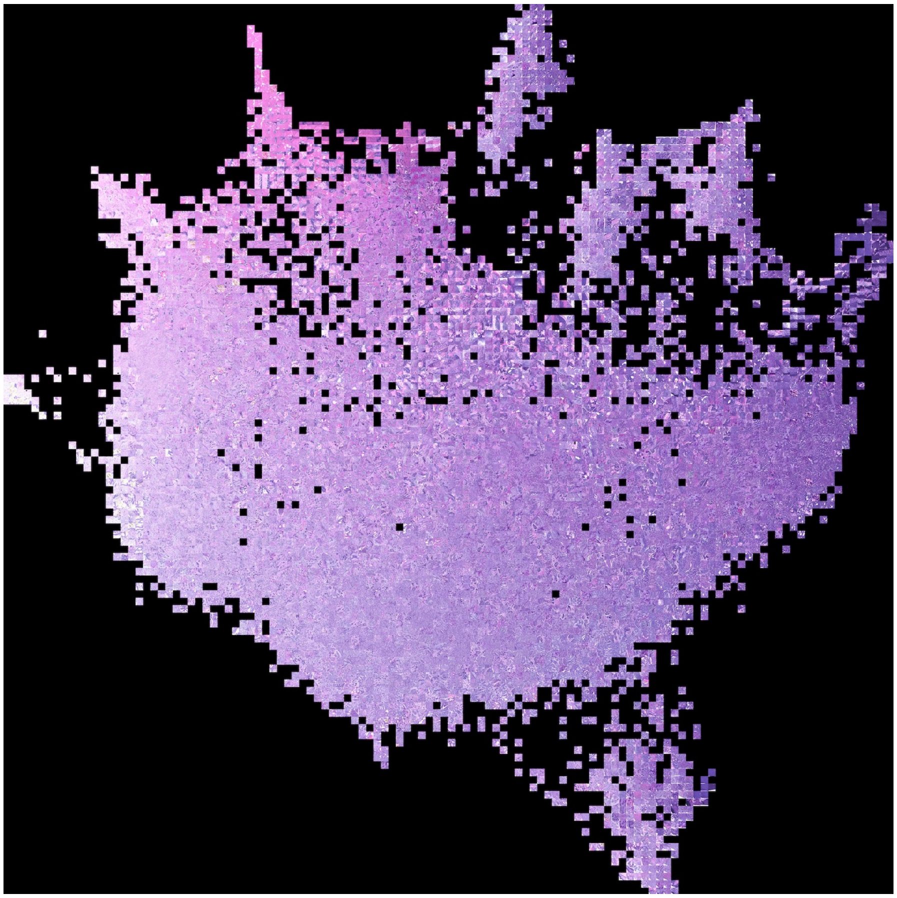

B

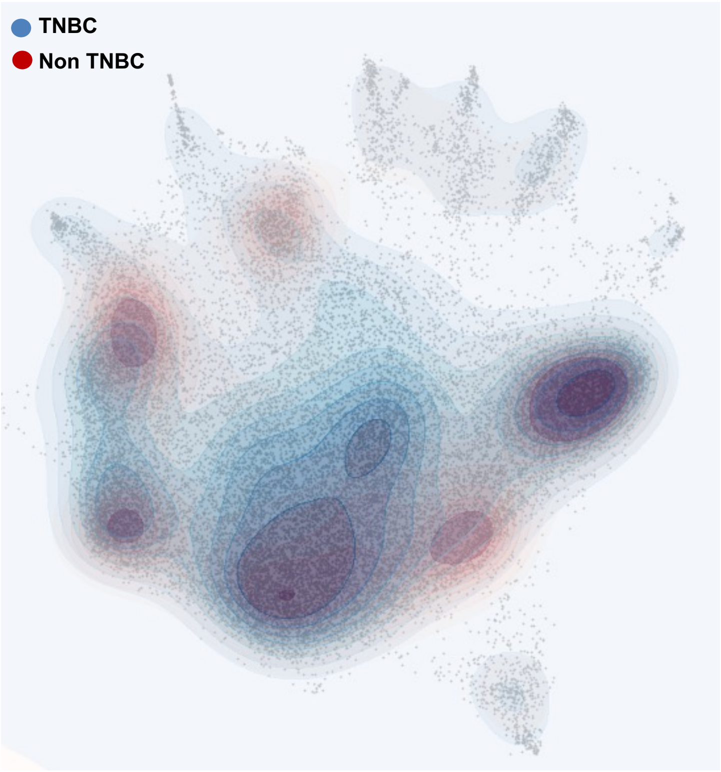

### Supplemental Figure S6

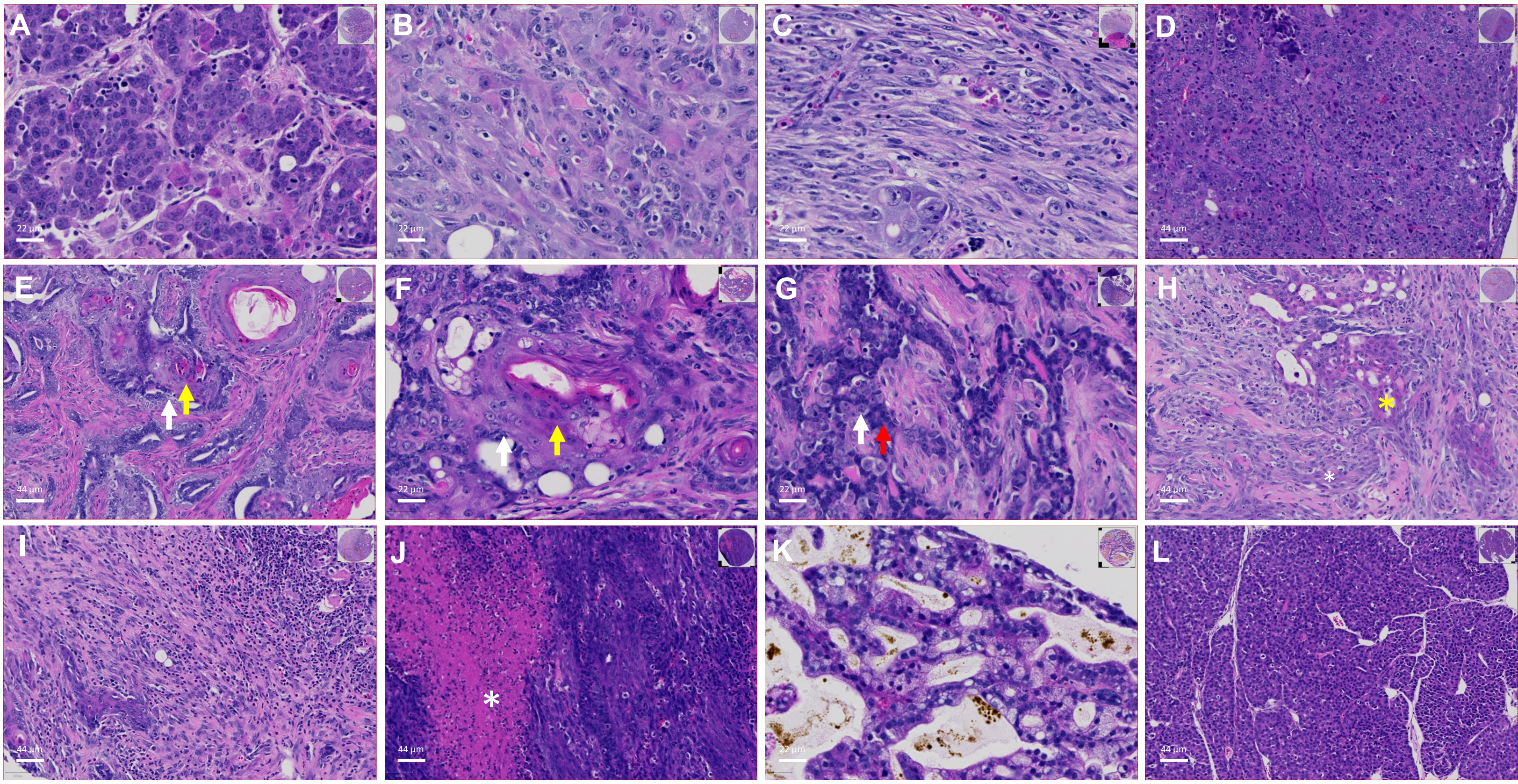


**M**

| Histologic characteristics of human TNBC | Stroma poor Mice | | Stroma rich Mice | | (N=80) | Frequency in mice TNBC model (%) |
|---|---|---|---|---|---|---|
| | N= 16 | Frequency (%) | N=64 | Frequency (%) | | |
| Low grade nuclear feature (A) | 8 | 50 | 9 | 14.0625 | 17 | 21.25 |
| High grade nuclear feature (B) | 6 | 37.5 | 8 | 12.5 | 14 | 17.5 |
| Sarcomatous transformation (C) | 2 | 12.5 | 8 | 12.5 | 10 | 12.5 |
| Solid growth pattern (D) | 8 | 50 | 12 | 18.75 | 20 | 25 |
| Squamoid metaplasia (E,F) | 0 | 0 | 38 | 59.375 | 38 | 47.5 |
| Myoepitheial proliferation (G,H) | 0 | 0 | 33 | 51.5625 | 33 | 41.25 |
| Geographic necrosis (J) | 1 | 6.25 | 16 | 25 | 17 | 21.25 |
| Fibrosis with stromal lymphocytic infiltrate (I) | 2 | 12.5 | 25 | 39.0625 | 27 | 33.75 |
| Clear cell change (K) | 0 | 0 | 9 | 14.0625 | 9 | 11.25 |
| Neuroendocrine differentiation (L) | 1 | 6.25 | 0 | 0 | 1 | 1.25 |

Column scaled
High
Low

**Supplemental Figure S7**

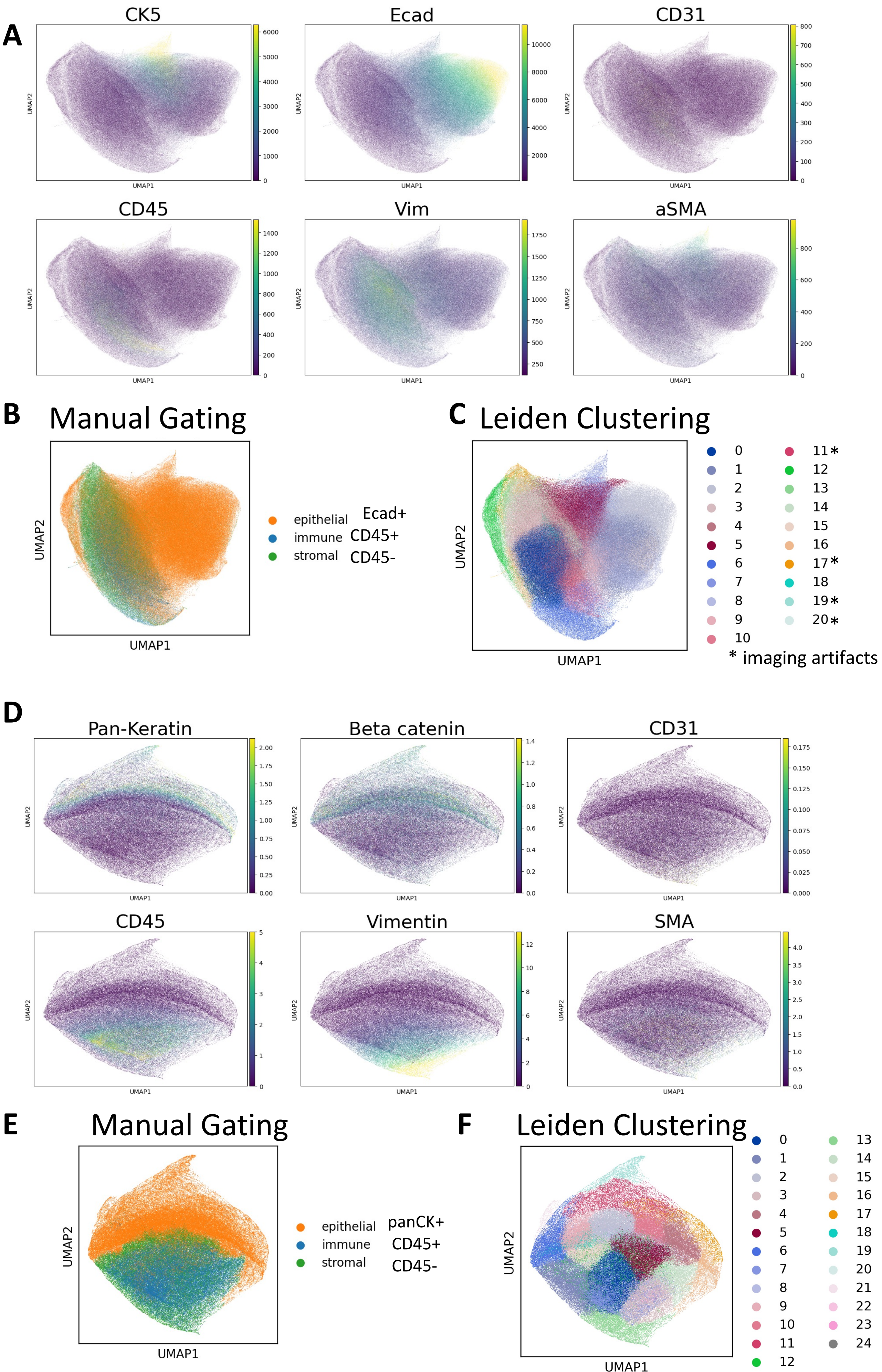

**Supplemental Figure S8**

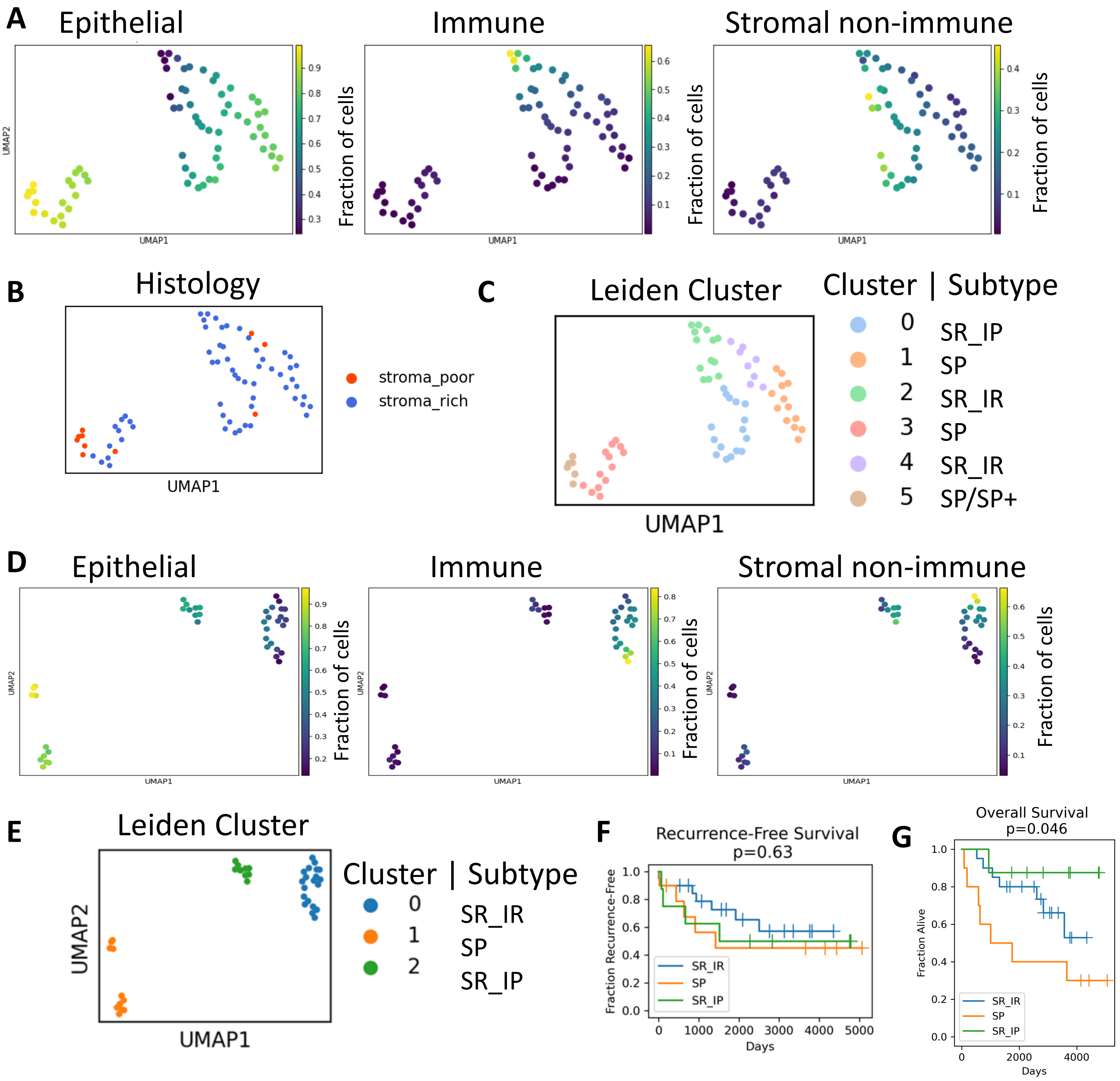

#### Supplemental Figure S9

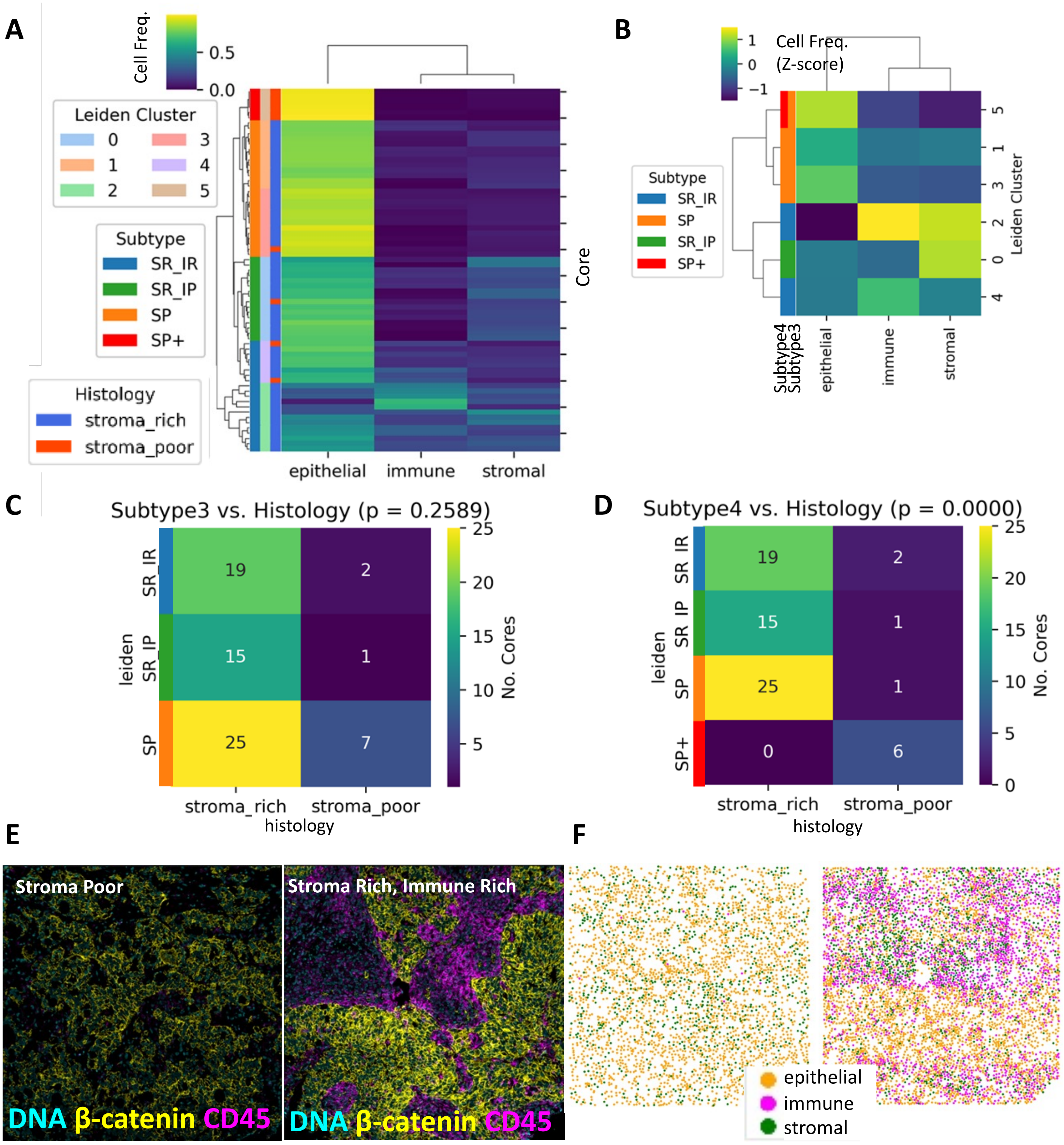

**Supplemental Figure S10**

**A** Mouse

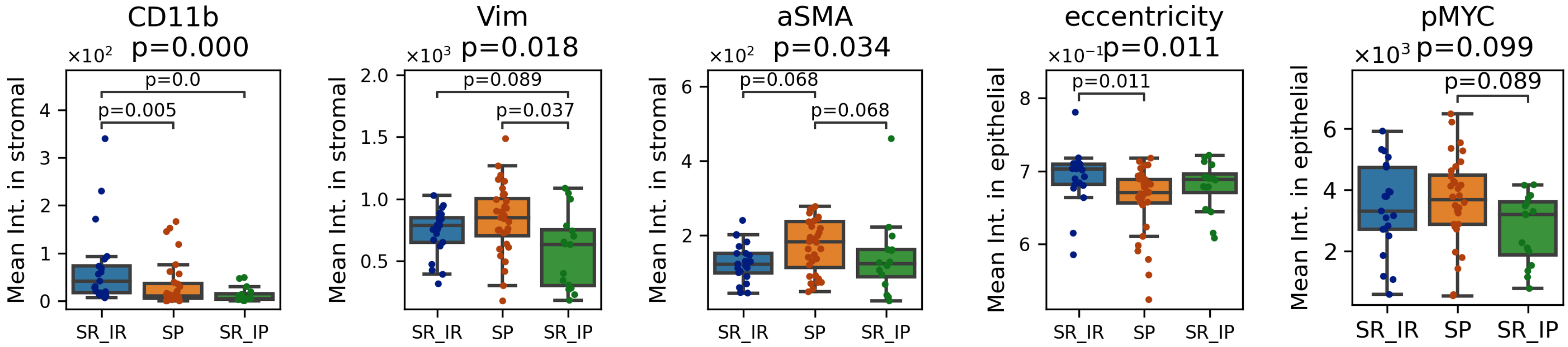


**B** Human

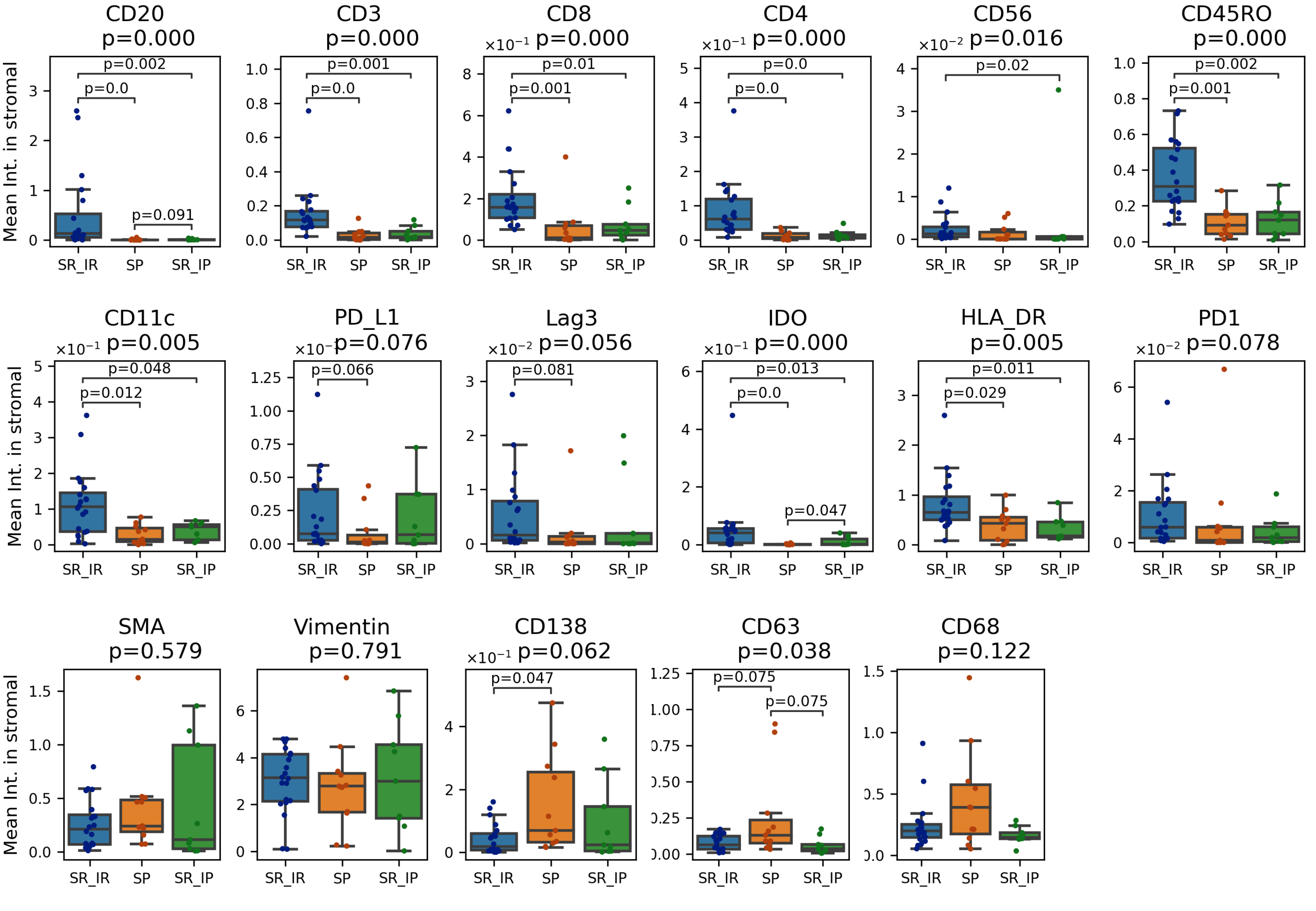


**C** Mouse Histology

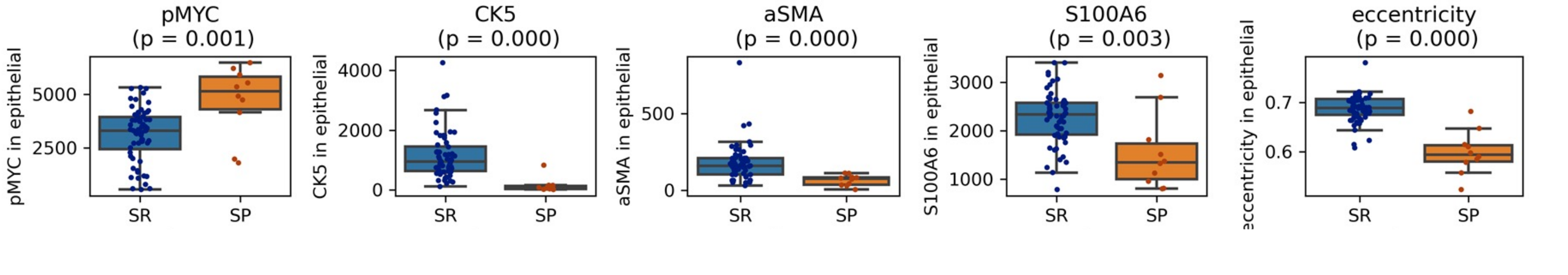

### Supplemental Figure S11

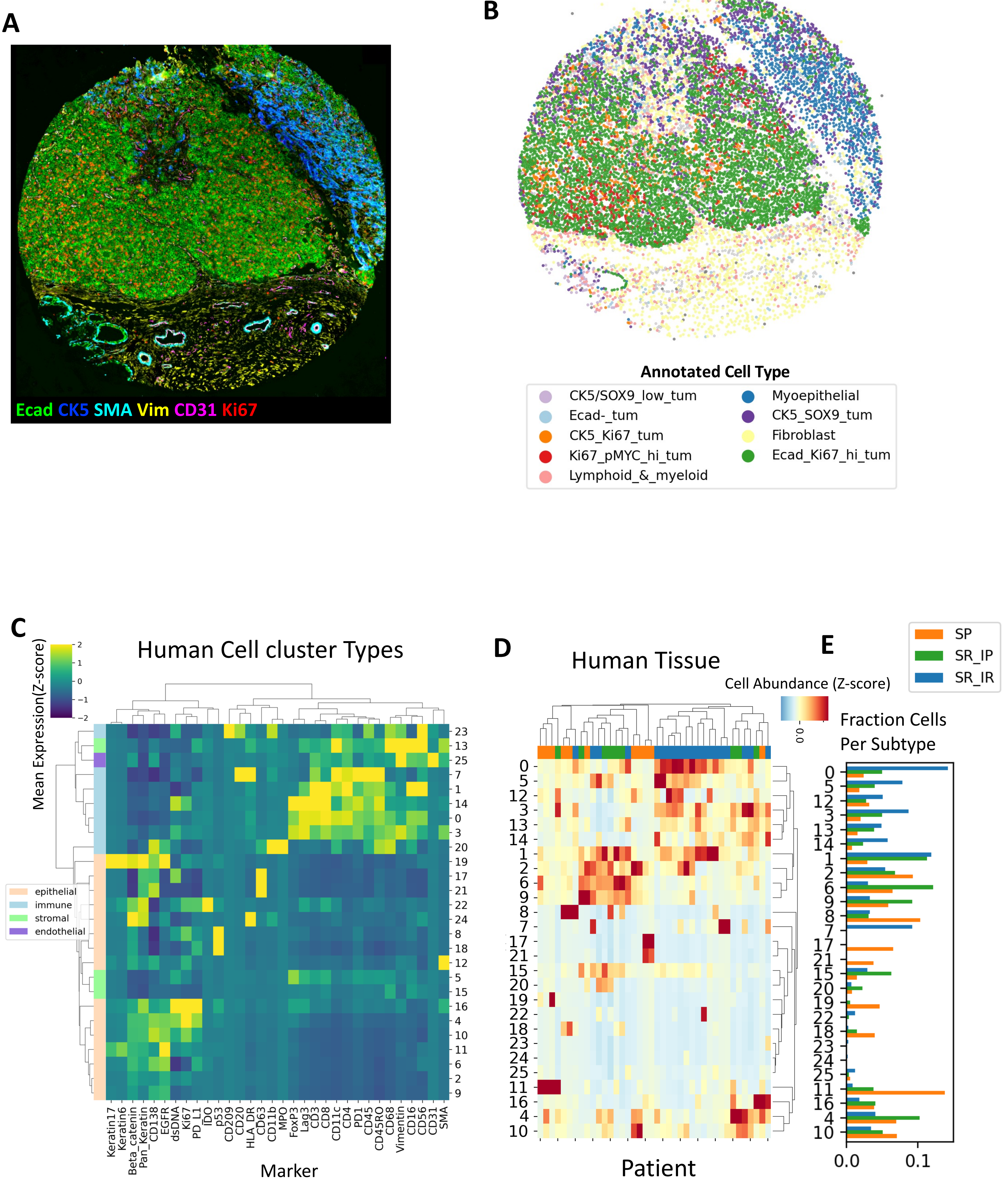

#### Supplemental Figure S12

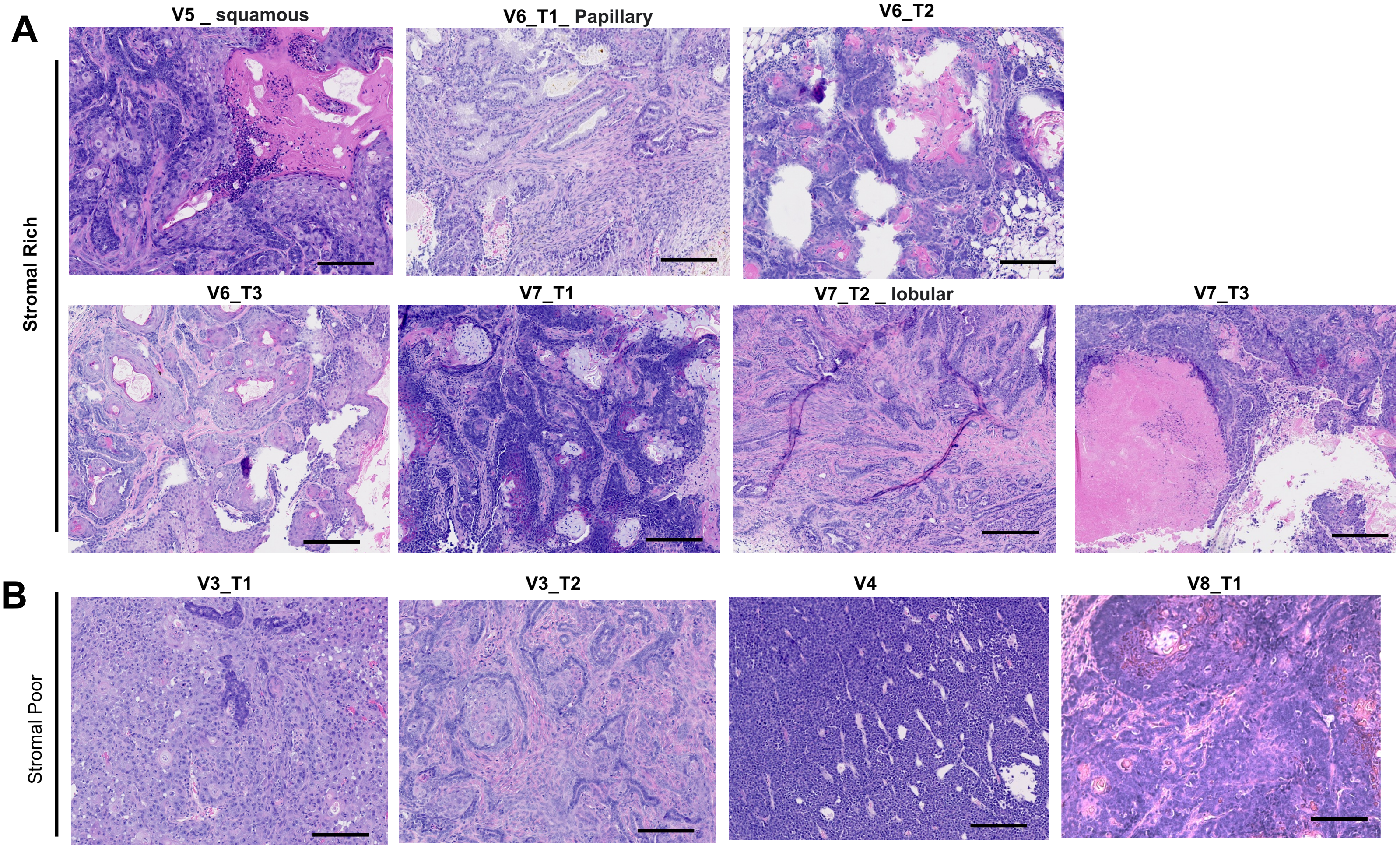

### Supplemental Figure S13

A

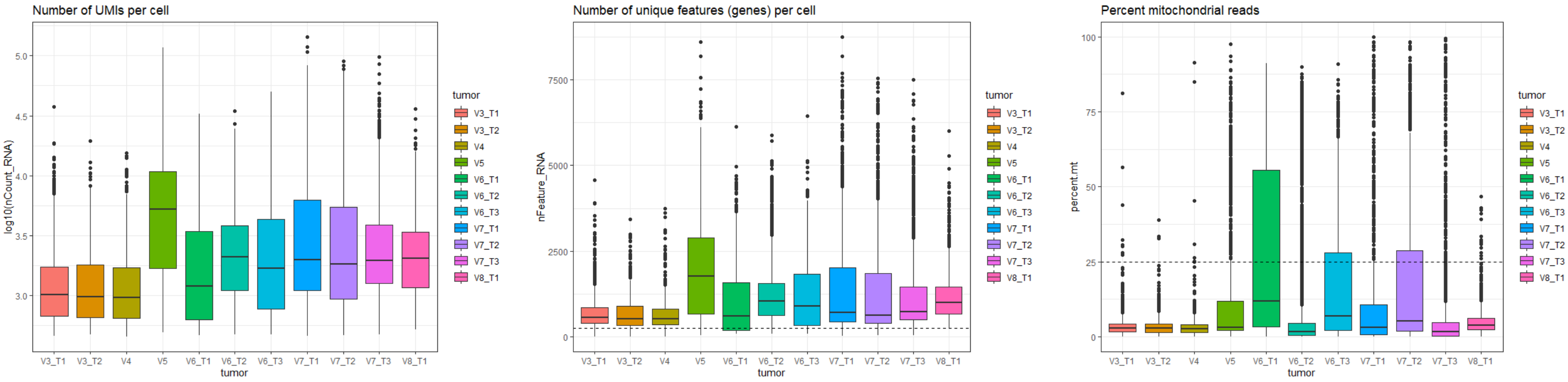


B.

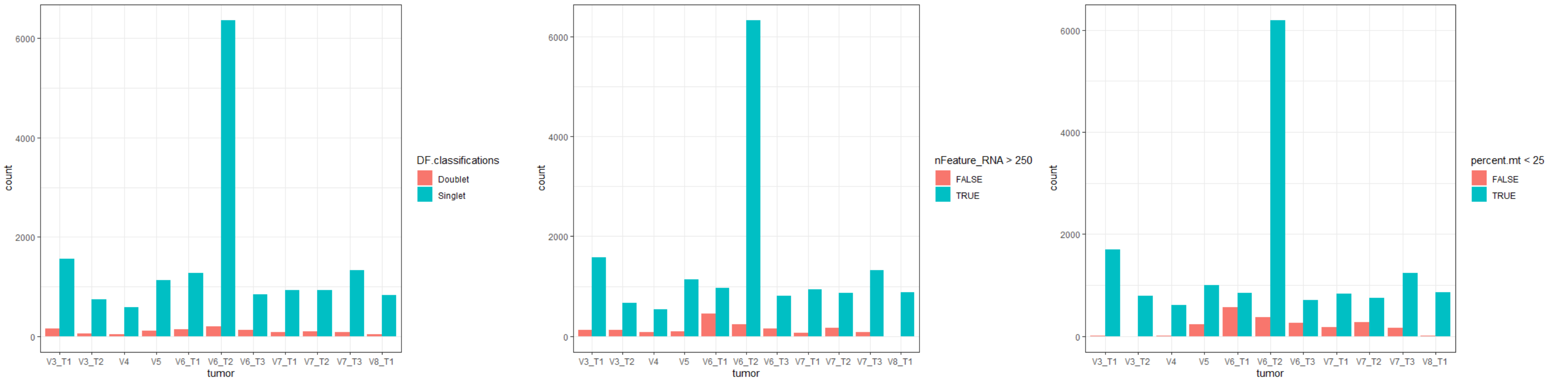


| tumor | nCells | mean_nCount_RNA | mean_nFeature_RNA | mean_percent.mt |
| :--- | ---: | ---: | ---: | ---: |
| V3_T1 | 1428 | 1385.333 | 672.6092 | 3.504835 |
| V3_T2 | 614 | 1467.952 | 691.5179 | 3.852925 |
| V4 | 490 | 1466.356 | 680.9755 | 3.584096 |
| V5 | 895 | 8792.437 | 2193.0559 | 4.367694 |
| V6_T1 | 700 | 3117.215 | 1291.2300 | 6.954764 |
| V6_T2 | 5995 | 2919.183 | 1200.7675 | 3.007906 |
| V6_T3 | 578 | 3331.370 | 1322.6003 | 6.579067 |
| V7_T1 | 742 | 5837.861 | 1437.7264 | 3.804922 |
| V7_T2 | 648 | 5179.649 | 1363.6250 | 5.043397 |
| V7_T3 | 1138 | 3969.579 | 1207.4446 | 2.544212 |
| V8_T1 | 814 | 2928.386 | 1170.1609 | 4.966005 |

**Supplemental Figure S14**

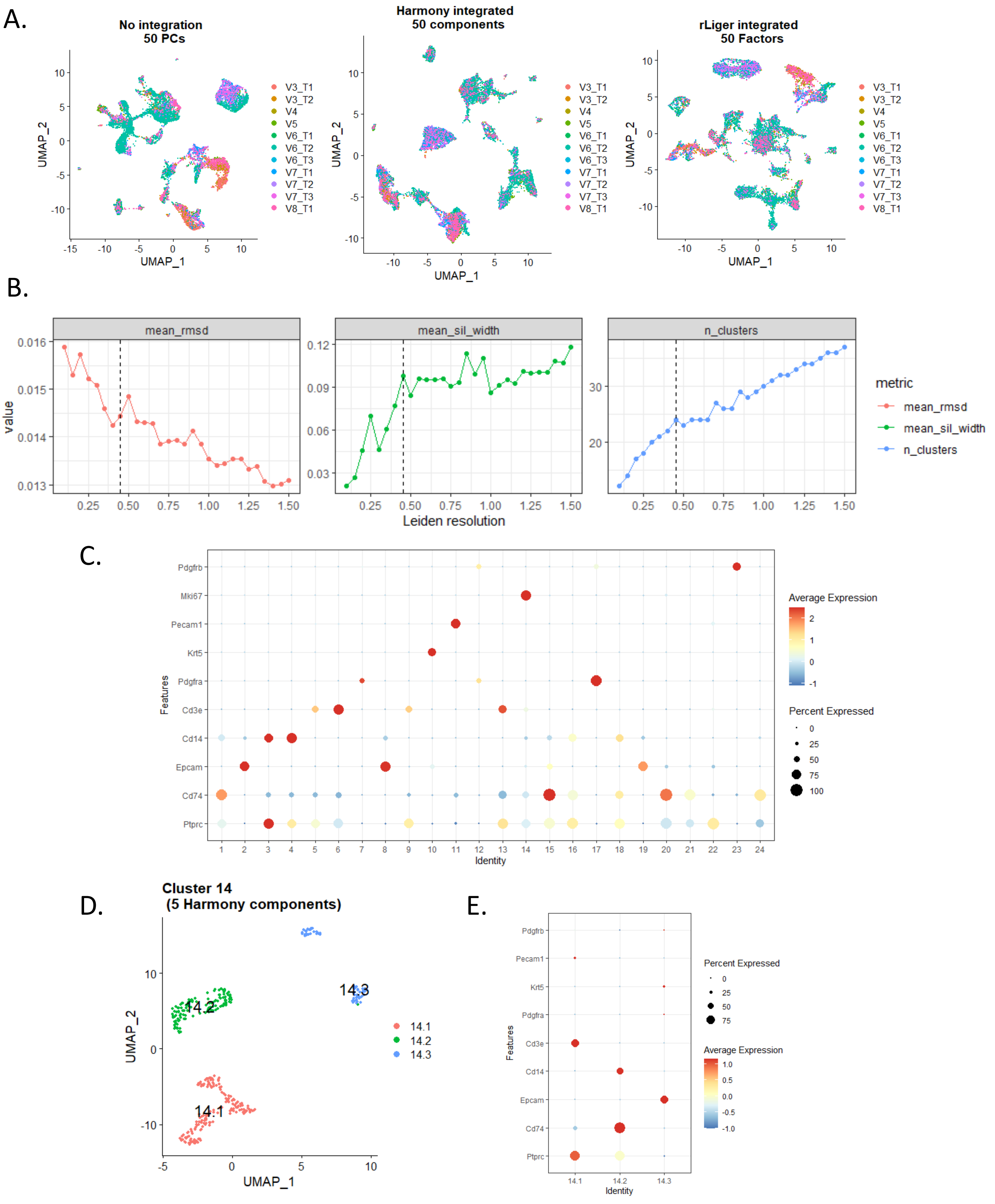

### Supplemental Figure S15

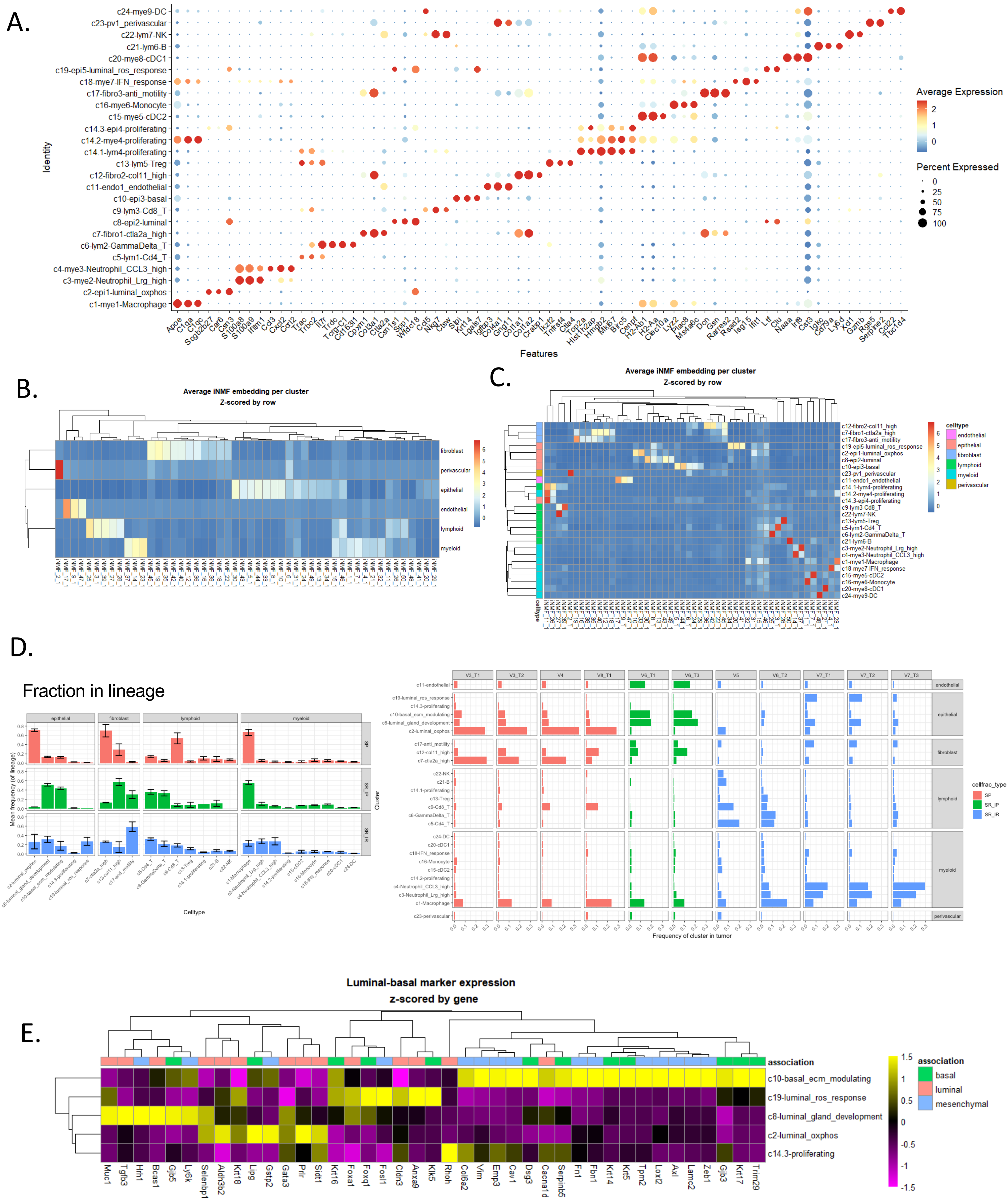

### Supplemental Figure S16

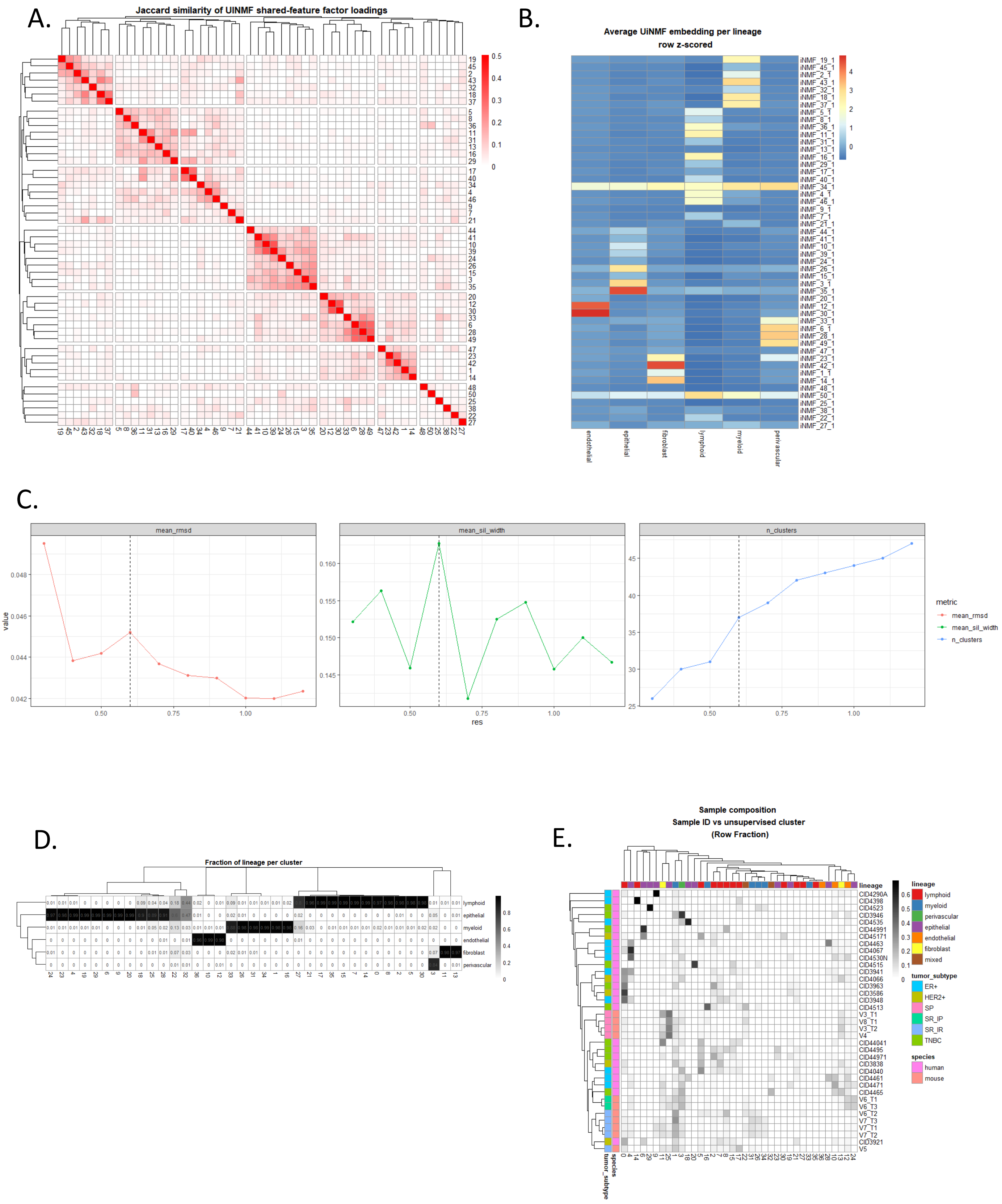

Supplemental Figure S17

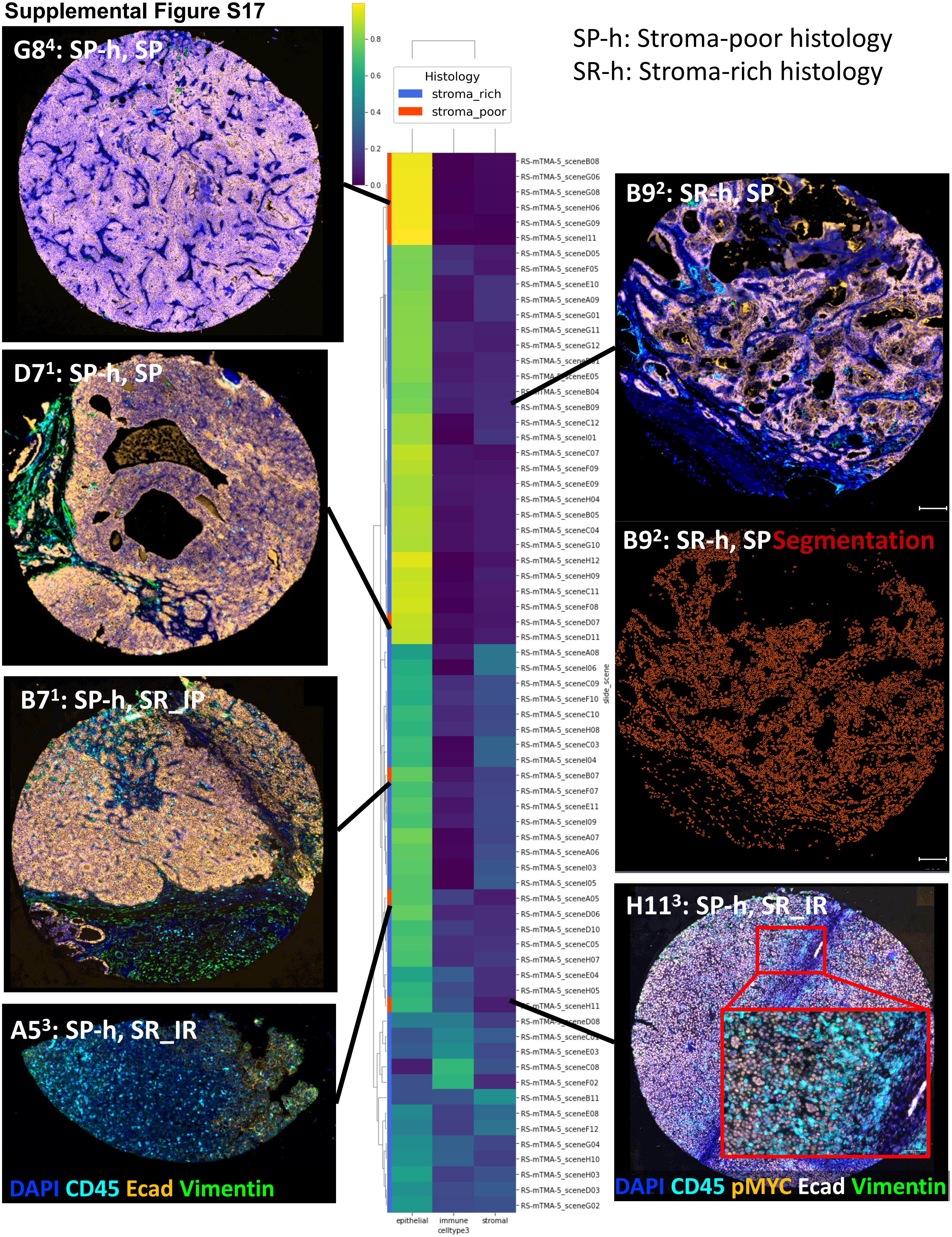
